# Identification of novel avian and mammalian deltaviruses provides new insights into deltavirus evolution

**DOI:** 10.1101/2020.08.30.274571

**Authors:** Masashi Iwamoto, Yukino Shibata, Junna Kawasaki, Shohei Kojima, Yung-Tsung Li, Shingo Iwami, Masamichi Muramatsu, Hui-Lin Wu, Kazuhiro Wada, Keizo Tomonaga, Koichi Watashi, Masayuki Horie

## Abstract

Hepatitis delta virus (HDV) is a satellite virus that requires hepadnavirus envelope proteins for its transmission. Although recent studies identified HDV-related deltaviruses in certain animals, the evolution of deltaviruses, such as the origin of HDV and the mechanism of its coevolution with its helper viruses, is unknown, mainly because of the phylogenetic gaps among deltaviruses. Here we identified novel deltaviruses of passerine birds, woodchucks, and white-tailed deer by extensive database searches and molecular surveillance. Phylogenetic and molecular epidemiological analyses suggest that HDV originated from mammalian deltaviruses and the past interspecies transmission of mammalian and passerine deltaviruses. Further, metaviromic and experimental analyses suggest that the satellite-helper relationship between HDV and hepadnavirus was established after the divergence of the HDV lineage from non-HDV mammalian deltaviruses. Our findings enhance our understanding of deltavirus evolution, diversity, and transmission, indicating the importance of further surveillance for deltaviruses.

## Introduction

Hepatitis delta virus (HDV) is the only member of the genus *Deltavirus* which is not assigned to a family (1). The HDV genome is an approximately 1.7-kb circular, negative single-stranded RNA, harboring a single open reading frame (ORF) encoding the small and large hepatitis delta antigens (S-HDAg, 24 kDa and L-HDAg, 27 kDa) that are translated from the same transcriptional unit via RNA-editing of the stop codon, which is catalyzed by the host protein ADAR1 (2–5). The 19 amino acid residue extension of the C-terminal region of L-HDAg contains a farnesylation site required to interact with helper virus envelope proteins (6). The genome structure of HDV is unique in that it has genomic and antigenomic ribozymes, which are essential for its replication (7, 8), and is highly self-complementarity, generating a rod-like structure (9–11). Although HDV can autonomously replicate, it requires an envelope protein of other “helper” viruses to produce infectious virions. Hepatitis B virus (HBV) (family *Hepadnaviridae*) provides the envelope proteins required for HDV transmission between humans (12). Approximately 15–20 million people worldwide are estimated to be infected with HDV among 350 million HBV carriers (13). Compared with mono-infection with HBV, coinfection of HDV and HBV accelerates the pathogenic effects of HBV, such as severe or fulminant hepatitis and progression to hepatocellular carcinoma, through unknown mechanisms (14).

The evolutionary origin of HDV presents an enigma. However, recent discoveries of deltaviruses of vertebrate and invertebrate species (15–18), significantly changed our understanding of deltavirus evolution. These non-HDV deltaviruses are distantly related to HDV but may share the same origin because of their similarly structured circular RNA genomes (approximately 1.7 kb), which encode DAg-like proteins, possess ribozymes sequences, and are highly self-complementary (15–18). These findings provide clues to the mechanism of deltavirus evolution. For example, a recent study hypothesizes that mammalian deltaviruses codiverged with their host mammalian species (18). However, the few known deltaviruses are highly divergent (15–18). Therefore, the phylogenetic gaps between the deltaviruses must be filled through the identification of putative novel deltaviruses.

The discoveries of non-HDV deltaviruses provides insights into the relationships between deltaviruses and their helper viruses. Recently identified non-HDV deltaviruses likely do not coinfect with hepadnaviruses, suggesting the presence of other helper viruses (15–18). This hypothesis is supported by absence of a large isoform of DAg, which is required for the interaction of HDV with the HBV envelope proteins, in rodent deltavirus (18). Further, viral envelope proteins of reptarenavirus and hartmanivirus, but not HBV, confer infectivity upon the snake deltavirus (19). These findings suggest that hepadnaviruses do not serve as helper viruses for non-HDV deltaviruses and that the deltavirus-hepadnavirus relationship is specific to the HDV lineage. However, the large phylogenetic gap between HDV and the few other deltaviruses makes it difficult to assess the hypothesis, raising the importance of further research.

In this study, to understand the evolution of deltaviruses, we analyzed publicly available transcriptome data and found novel mammalian and avian deltaviruses. Our phylogenetic analysis suggests that HDVs originated from non-HDV mammalian deltaviruses and does not support the codiversification hypothesis of deltavirus and mammalian evolution. Moreover, *in silico* and experimental analyses, together with previous findings, suggest that the satellite-helper relationship between HDV and hepadnavirus was established after the divergence of the novel non-HDV mammalian deltaviruses and the HDV lineage. Further, we present evidence for recent interfamily transmission of deltaviruses among passerine birds. Our findings therefore provide novel insights into the evolution of deltaviruses.

## Results

### Identification of deltavirus-related sequences in avian and mammalian transcriptomes

We first assembled 46,359 RNA-seq data of birds and mammals. Using the resultant contigs as queries, we identified five deltavirus-related contigs in the SRA data of birds and mammals, including the zebra finch (*Taeniopygia guttata*), common canary (*Serinus canaria*), Gouldian finch (*Erythrura gouldiae*), Eastern woodchuck (*Marmota monax*), and white-tailed deer (*Odocoileus virginianus*). We named the deltavirus-like sequences Taeniopygia guttata deltavirus (tgDeV), Serinus canaria-associated deltavirus (scDeV), Erythrura gouldiae deltavirus (egDeV), Marmota monax deltavirus (mmDeV), and Odocoileus virginianus (ovDeV), respectively (Table 1).

**Table 1.** Summary of novel deltaviruses

| Virus name |  | Host species | Tissue | SRA | DDBJ | Contig<br>length (nt) | GC content<br>(%) | BLASTx best hit |  |  |
| --- | --- | --- | --- | --- | --- | --- | --- | --- | --- | --- |
|  |  |  |  | accession | accession |  |  | Virus name | Accession | Identity (%) |
| Taeniopygia guttata DeV | tgDeV | *Taeniopygia guttata* | Scapulohumeralis<br>caudalis | SRR2545946 | BR001665 | 1706 | 56.6 | Rodent deltavirus | QJD13558 | 63.3 |
| Marmota monax DeV | mmDeV | *Marmota monax* | Liver | SRR2136906 | BR001661 | 1712 | 53.4 | Hepatitis delta virus | AIR77039 | 60.0 |
| Odocoileus virginianus DeV | ovDeV | *Odocoileus virginianus* | Pedicle | SRR4256033 | BR001662 | 1690 | 56.4 | Hepatitis delta virus | AHB60712 | 66.7 |
| Erythrura gouldiae DeV | egDeV | *Erythrura gouldiae* | Skin | SRR7504989 | BR001660 | 596 | 59.4<sup>a)</sup> | Rodent deltavirus | QJD13555 | 63.5 |
| Serinus canaria-associated DeV | scDeV | *Serinus canaria* | Skin | SRR2915371 | BR001664 | 761 | 54.4<sup>a)</sup> | Hepatitis delta virus | AIR77012 | 36.0 |
| Pardaliparus venustulus DeV | pvDeV | *Pardaliparus venustulus* | Lung, Kidney,<br>Cardiac muscle,<br>Flight muscle,<br>Liver | SRR7244693<br>SRR7244695<br>SRR7244696<br>SRR7244697<br>SRR7244698 | BR001663 | 1708 | 55.8 | Rodent deltavirus | QJD13562 | 62.4 |
| Lonchura striata DeV | lsDeV | *Lonchura striata* var.<br>*domestica* | Blood | - | LC575944 | 1708 | 56.2 | Rodent deltavirus | QJD13555 | 62.9 |

a) GC content of the partial genome sequences.

The amino acid sequences tlanslated from the contigs are 36.0%–66.7% identical to those of the DAg proteins of known deltaviruses (Table 1, Supp Tables 1 and 2). The tgDeV, mmDeV, and ovDeV contigs, which comprise approximately 1,700 nucleotides, encode one ORF with a sequence similar to those of DAg genes of known deltaviruses (Fig 1a, Table 1, Supp Tables 1 and 2). In contrast, the contigs scDeV and egDeV are 761 and 596 nucleotides in length, respectively (Fig. 1b, Supp Tables 1 and 2). Note that the nucleotide sequences of tgDeV and egDeV are 97.7% identical, and we therefore analyzed tgDeV instead of tgDeV and egDeV.

**Figure 1.**
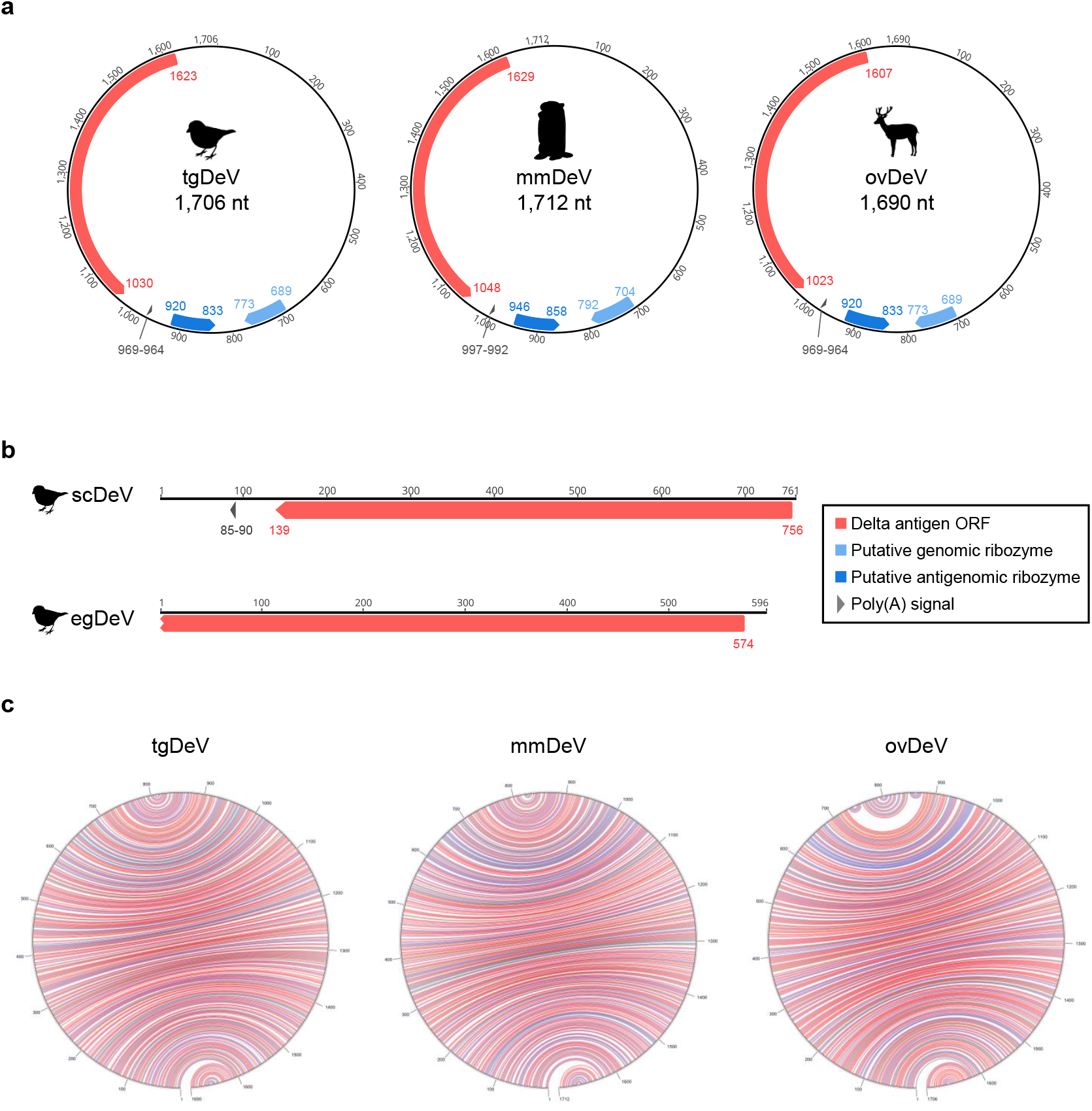
Genome organization of novel deltaviruses. Genomes of **(a)** tgDeV, mmDeV, and ovDeV (complete genomes) and **(b)** scDeV and egDeV (partial genomes). Annotations (ORF, poly-A signal, and ribozymes) are shown by colored arrow pentagons. The numbers indicate nucleotide positions. **(c)** Self-complementarities of novel deltaviruses. The predicted RNA structures were visualized using the Mfold web server (58). Red, blue, and green arcs indicate G-C, A-U, and G-U pairs, respectively.

### Genome structures of novel avian and mammalian deltaviruses

The three contigs (tgDeV, mmDeV, and ovDeV) are almost identical in length to the full-length genomes of known deltaviruses. We therefore checked for potential circularity of the contigs. Dot-plot analyses revealed that each of both ends of these three contigs is identical (Supp Fig. 1), suggesting that the contigs were derived from circular RNAs. We further mapped the original RNA-seq data to the corresponding circularized contigs using the Geneious mapper, revealing that some of the reads properly spanned the junctions (data not shown), indicating that these contigs are derived from circular RNAs. Therefore, we designated the resultant circular contigs of tgDeV, mmDeV, and ovDeV as full-length novel deltavirus genomes (1,706, 1,712, and 1,690 nucleotides, respectively) (Fig. 1a). These novel genomes are characterized by high self-complementarity, genomic and antigenomic ribozymes, and poly(A) signals, which are conserved among known deltaviral genomes (Fig. 1 and Supp Table 2) (15–18). Further, the predicted secondary structures of the ribozymes are highly similar to those of HDV as well as those of other deltaviruses (Supp Fig. 2).

### Characterization of DAg proteins encoded by the novel deltaviruses

We next characterized the putative DAg proteins encoded by the novel deltaviruses. Most of their biochemical features, biologically relevant amino acid residues, and functional domains (15–18) are conserved among the DAg proteins (Fig. 2a). The isoelectric points of DAg proteins from the novel deltaviruses range from 10.35 to 10.63 (Supp Table 2), which are nearly identical to those of known deltaviruses. All the post-translational modification sites in HDAg are conserved among those of all DAg proteins of the novel deltaviruses, except for the serine phosphorylation site on scDeV-DAg (Fig. 2a). The NLS is conserved among the DAg proteins, although location of the predicted NLS of scDeV DAg protein differs (Fig. 2a).

**Figure 2.**
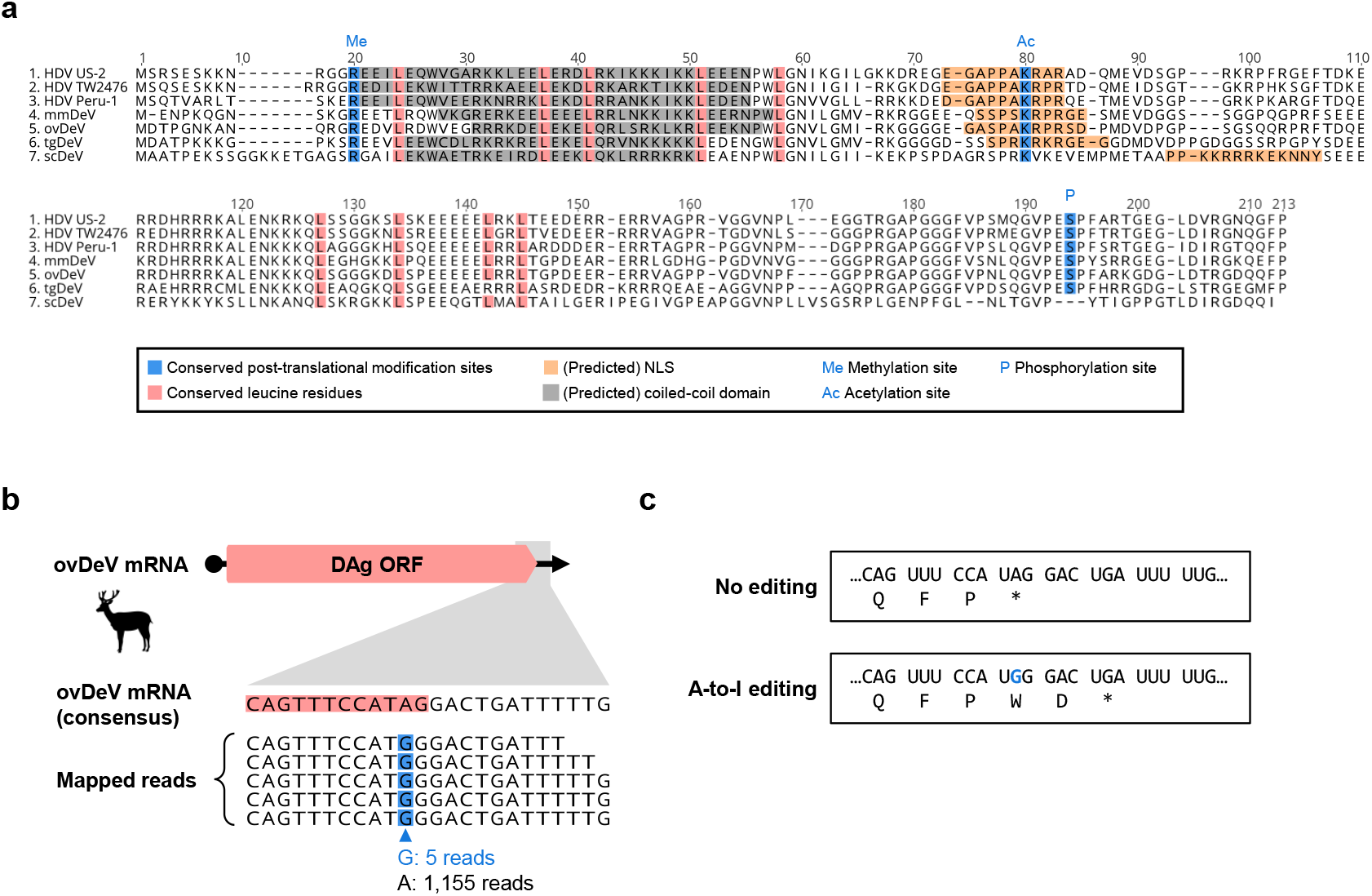
Amino acid sequence characterization of putative delta antigens of novel deltaviruses. **(a)** Alignment and functional features of the putative S-HDAg and DAgs of representative HDVs and novel deltaviruses. (Putative) functional domains are shown by colored boxes. Me: arginine methylation site, Ac: lysine acetylation site, P: Serine phosphorylation site. **(b)** ovDeV mRNA (upper panel) and a possible A-to-I RNA-editing site (lower panel). Consensus ovDeV-DAg mRNA sequence and mapped read sequence with potential RNA-edited nucleotides (blue boxes). Pink boxes indicate the ORF of ovDeV DAg. **(c)** Deduced amino acid sequences of ovDeV-DAg proteins translated from the viral mRNA with or without RNA-editing. The blue letter shows the possible RNA-editing site.

We next investigated whether the novel deltaviruses utilize A-to-I RNA-editing. To answer this question, we mapped short reads of the SRA data, which we initially used to detect the deltaviruses, to identify the nucleotide variations among the stop codons. We found a potential RNA-editing site within the stop codon of the ovDeV-DAg gene, in which there was 0.4% nucleotide variation (5 of 1160 reads) at the second nucleotide position of the stop codon (UAG), all of which were G instead of the consensus nucleotide A (Fig. 2b). The quality scores of the five G variants ranged from 35 to 41 (Supp Fig. 3), which likely exclude the possibility of a sequencing error. This variation may be explained by A-to-I editing by ADAR1, as known for HDV (2). However, possible RNA-editing generates a protein two amino acid residues longer because of a stop codon immediately downstream (Fig. 2c). Further, the C-terminal farnesylation motif (CXXQ) required for the interaction with hepadnaviral envelope proteins (6) was absent from the longer product. These observations suggest that even if RNA-editing occurs, the resultant gene product does not contribute to the interaction with hepadnaviral envelope proteins. Further, we were unable to identify nucleotide variations of the mapped reads at the stop codons in the genomes of tgDeV and mmDeV (data not shown).

### The novel deltaviruses potentially replicate in their hosts

To determine whether the novel deltaviruses potentially replicate in their respective, putative host species, we evaluated the mapping pattern of viral reads described above. We found that the read depths of the predicted transcribed regions (the DAg coding regions to poly-A signals) were much greater than those of the other genomic regions (Fig. 3), indicating that most viral reads were derived from viral mRNAs. These findings suggest that the novel deltaviruses replicate in their hosts.

**Figure 3.**
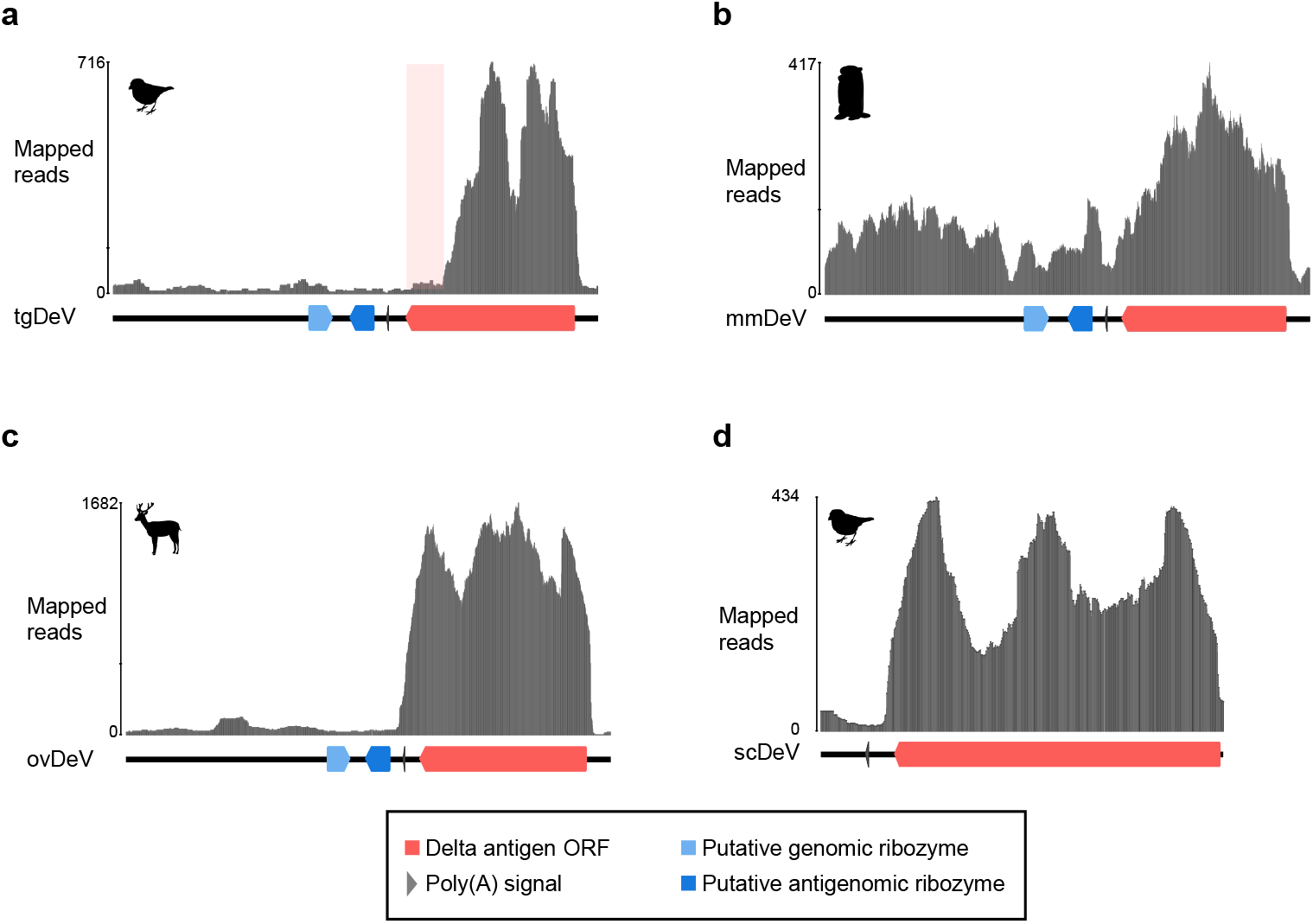
Mapping coverages of original short reads of each contig. Mapped read graphs of **(a)** tgDeV, **(b)** mmDeV, and **(c)** ovDeV. Lines, arrow pentagons, and arrowheads indicate viral genomes, ribozymes, and poly(A) signals, respectively. The numbers above the graphs show nucleotide positions. The light pink box indicates a low read depth region in the putative transcript of tgDeV.

The mapping pattern on tgDeV differed slightly from the others. Specifically, although the read depth of the DAg ORF region was higher, the reads represented only 80% of the ORF (Fig. 3a). This trend was apparent in another tgDeV-positive RNA-seq data (Supp Fig. 4). However, it is not clear whether this is attributed to an artifact or actually reflects the transcription pattern of tgDeV.

### Transmission of tgDeV- and tgDeV-like viruses among passerine birds

We next investigated whether the novel deltaviruses are transmitted among animal populations. We first analyzed tgDeV infections in birds using RNA-seq data (Table 2 and Supp Table 3). Among 6453 SRA data, tgDeV-derived reads were identified in 34 SRAs, including the SRAs in which tgDeV and egDeV were initially detected. The 34 tgDeV-positive SRA data were obtained from tissues such as blood, kidney, and muscles, suggesting broad tropism and viremia, or systemic infection, or both, with tgDeV.

**Table 2.** Detection of deltavirus-derived reads in RNA-seq data.

| Virus | BioProject/<br>BioStudy | SRA | Host |  | RPM <sup>a)</sup><br>(read per million) | Tissue |
| --- | --- | --- | --- | --- | --- | --- |
|  |  |  | Taxonomy |  |  |  |
|  |  |  | Family | Species |  |  |
| tgDeV | PRJNA297576 | SRR2545943 | Estrildidae | *Taeniopygia guttata* | 10.28 | Pectoralis |
|  |  | SRR2545944 |  |  | 1.02 | Scapulohumeralis caudalis |
|  |  | SRR2545946 |  |  | 56.73 | Scapulohumeralis caudalis |
|  | PRJNA558524 | SRR9899549 | Emberizidae <sup>b)</sup> | *Emberiza melanocephala* | 3.11 | Blood |
|  | PRJNA470787 | SRR7244693 | Paridae | *Pardaliparus venustulus* | 10.68 | Lung |
|  |  | SRR7244695 |  |  | 2.07 | Kidney |
|  |  | SRR7244696 |  |  | 2.65 | Cardiac muscle |
|  |  | SRR7244697 |  |  | 7.12 | Flight muscle |
|  |  | SRR7244698 |  |  | 1.77 | Liver |
|  | PRJNA478907 | SRR7504989 | Estrildidae | *Erythrura gouldiae* | 1.07 | Skin |
| mmDeV | PRJNA291589 | SRR2136906 |  | *Marmota monax* | 70.86 | Liver |
|  |  | SRR2136907 |  |  | 63.08 | Liver |
|  |  | SRR2136916 |  |  | 1.02 | Liver |
|  | SRP011132 | SRR437934 |  |  | 46.34 | PBMC |
|  |  | SRR437938 |  |  | 19.83 | PBMC |
| ovDeV | PRJNA317745 | SRR4256033 |  | *Odocoileus virginianus* | 180.73 | Pedicle |
| scDeV | PRJNA300534 | SRR2915371 |  | *Serinus canaria* | 9.79 | Skin |

The full version of the table is available as Supplementary Table 3.
a) This table only shows the samples with RPM >1.
b) Emberizidae is regarded as the subfamily Emberizinae of the family Fringillidae in TimeTree.

Further, tgDeV sequences were detected in several bird species such as the black-headed bunting (*Emberiza melanocephala*) and yellow-bellied tit (*Pardaliparus venustulus*). All tgDeV-positive bird species belong to the order Passeriformes. These tgDeV-positive SRA data are included in the nine BioProjects deposited by independent researchers, and thus the birds were likely from different sources. Further, the tgDeV-positive sample in SRR9899549 (BioSample accession: SAMN12493457) is derived from a black-headed bunting caught in the wild. These data suggest that tgDeV (or tgDeV-like viruses) circulate among diverse passerine birds, even in the wild.

During the above analysis, we found that SRA data from the yellow-bellied tit (SRR7244693 and SRR7244695–SRR7244698) contain many reads mapped to the tgDeV genome. Therefore, we extracted the mapped reads of SRA data and performed *de novo* assembly. We obtained a 1707-nt circular complete genome sequence, which we designated pvDeV The pvDeV nucleotide sequence is 98.2% identical to that of tgDeV, and the properties of its DAg protein sequence are similar to those of tgDeV DAg (Supp Fig. 5).

We next employed real-time RT-PCR to further evaluate potential deltavirus infections of passerine birds. We analyzed 30 and 5 whole-blood samples from zebra and Bengalese finches *(Lonchura striata* var. *domestica*), respectively, and found that one Bengalese finch was positive for real-time RT-PCR test. To exclude the possibility of contamination of a plasmid used as a control for real-time PCR, we performed RT-PCR using a primer set that distinguishes viral from plasmid amplicons (Figs. 4a and b). We obtained a band of the expected size only from the cDNA sample (Fig. 4c), revealing that the bird was truly positive for a tgDeV-like virus. Therefore, we named this virus lsDeV, and further analysis revealed that its full-length genome nucleotide sequence (1708 nt) is 98.2% and 98.4% identical to those of tgDeV and pvDeV, respectively. Moreover, its genome features are almost identical to those of tgDeV and pvDeV (Supp Fig. 5).

**Figure 4.**
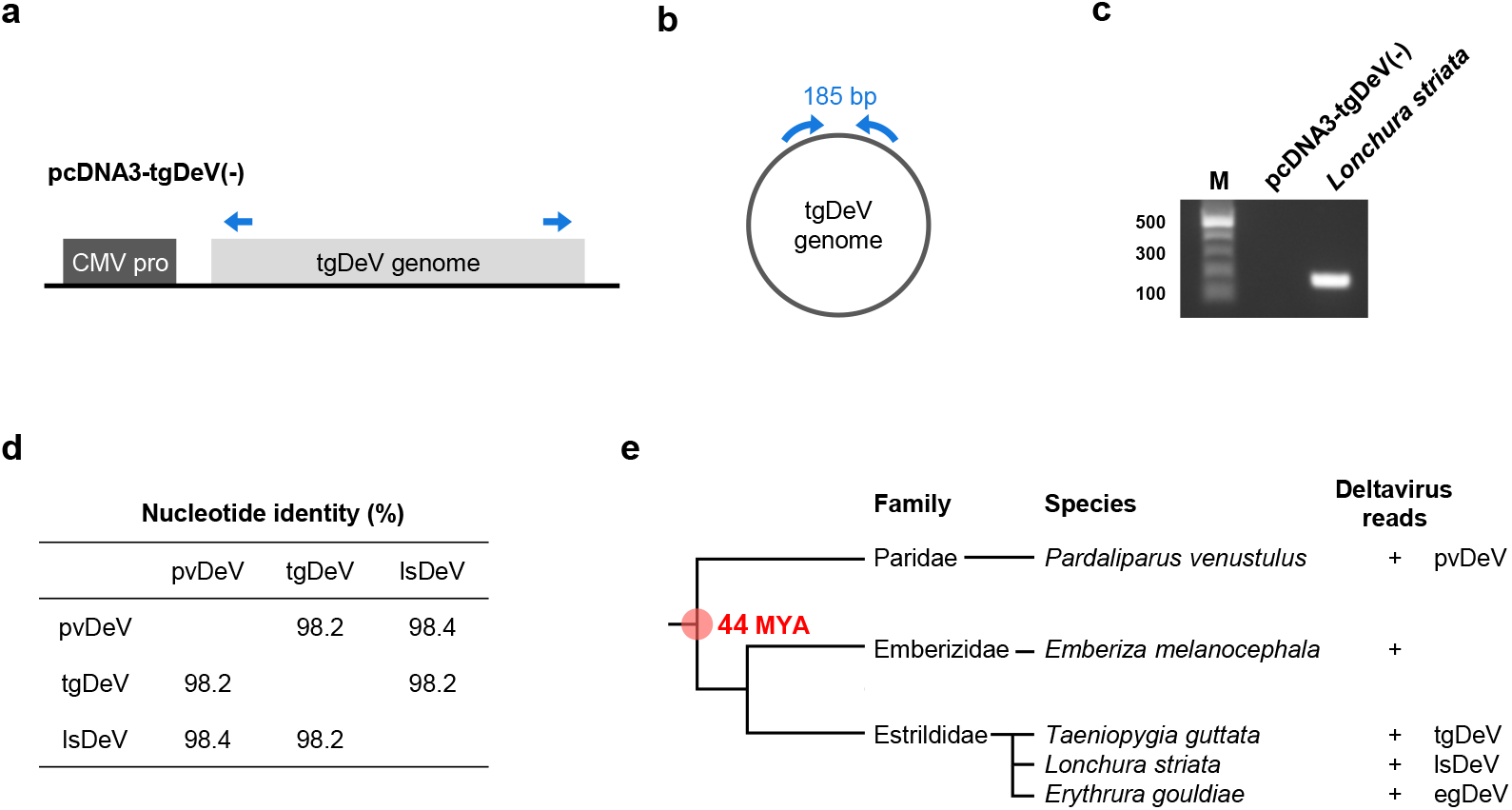
Interfamily transmission of deltaviruses among passerine birds. **(a–c)** RT-PCR detection of a deltavirus from *Lonchura striata.* **(a)** Plasmid used for the establishment of real-time PCR detection system for tgDeV and **(b)** the tgDeV circular genome. The blue arrows indicate the primers used for endo-point RT-PCR detection. **(c)** Endo-point RT-PCR for detection of the circular deltavirus genome. M, 100-bp ladder marker. **(d)** Pairwise nucleotide identities between deltaviruses detected in passerine birds. **(e)** Phylogenetic tree of passerine birds positive for deltaviruses. Phylogenetic tree of birds and deltavirus infections are indicated. MYA: million years ago.

### Evidence for recent interfamily transmission of deltaviruses among passerine birds

We found that the sequence similarities among the passerine deltaviruses (tgDeV, pvDeV, and ls DeV) (Fig. 4d) were not consistent with evolutionary codivergence. According to the TimeTree (20), deltavirus–positive passerine birds diverged approximately 44 million years ago (Fig. 4e and Supp Fig. 6). Considering the rapid evolutionary rates of HDVs (approximately 10^−3^ substitutions per site per year) (21–23), it is unlikely that these viruses codiverged with their hosts. Most likely, interfamily transmission occurred relatively recently among passerine birds.

### Transmission of mmDeV among woodchucks

We similarly analyzed mmDeV infections using SRA data for the order Rodentia, other than mice (*Mus musculus*) and rats (*Rattus norvegicus*). Our analysis of 4776 SRA datasets detected mmDeV reads in 20 SRA data of seven woodchucks (Table 2 and Supp Table 3). Although these mmDeV-positive SRA data were contributed by the same research group, the animals were apparently obtained at different times (24, 25), suggesting that mmDeV was transmitted among woodchucks. The mmDeV-positive SRA data are derived from samples of liver or peripheral blood mononuclear cells.

We next used real-time RT-PCR to analyze 81 woodchuck samples (liver, n= 43; serum, n = 38). However, mmDeV was undetectable (data not shown), which may be explained by the absence of mmDeV infection or clearance, low level of infection, or both.

### No evidence of transmission of other deltaviruses

We next focused on ovDeV and scDeV sequences of ruminant animals and passerine birds, respectively (Table 2 and Supp Table 3). We detected ovDeV-derived reads only in five SRA data. The SRA data were obtained from brain, muscle, testis, pedicles, and antlers, suggesting systemic infection, viremia, or both. However, we were unable to determine if these samples were derived from multiple individuals. We detected scDeV-derived reads only from the SRA data in which the virus was initially detected. We therefore were unable to provide evidence for the transmission of ovDeV and scDeV in their host animals.

### Phylogenetic relationships among deltaviruses

To decipher the evolutionary relationships among deltaviruses, we conducted a phylogenetic analysis using known and the novel deltavirus sequences discovered here. We did not include sequences of recently identified fish, toad, newt, termite, and duck-associated deltaviruses because they share very low amino acid sequence identities with the novel deltaviruses as well as with HDVs (Fig. 5a), which may reduce the accuracy of tree (16). We further excluded scDeV for this reason, and we did not include tgDeV-like viruses, because their sequences are nearly identical to that of tgDeV The reconstructed tree shows that the newly identified tgDeV forms a strongly supported cluster with snake DeV and rodent DeV, although they are distantly related to each other (Fig. 5b). Note that mmDeV and ovDeV are more closely related to HDVs than the other deltaviruses.

**Figure 5.**
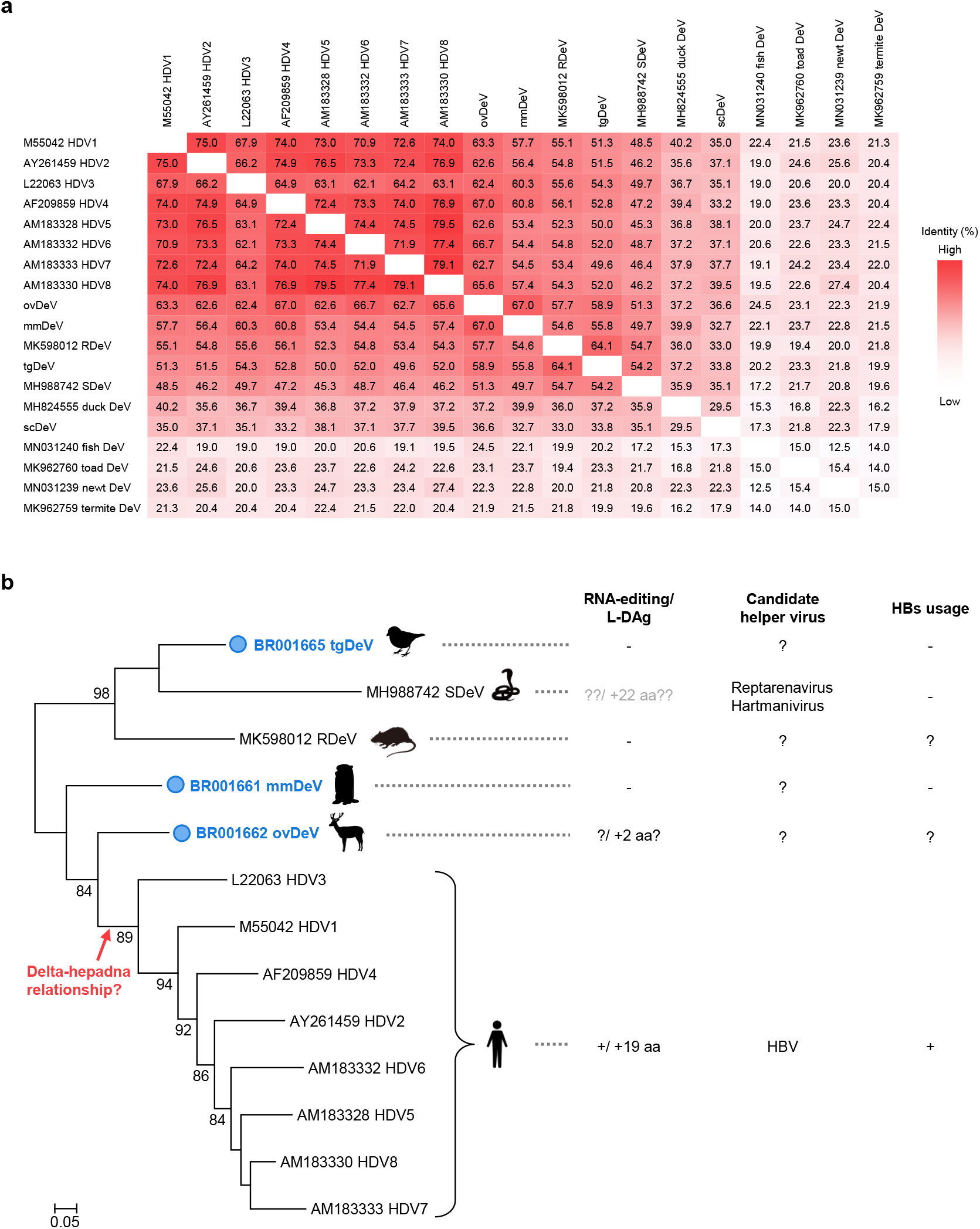
Phylogenetic analysis of deltaviruses. **(a)** Heat map of pairwise amino acid sequence identities between deltaviruses. **(b)** The phylogenetic tree was inferred by the maximum likelihood method using an amino acid sequence alignment of representative deltaviruses. Known phenotypes (RNA-editing and expression of the large isoform of DAg protein) and helper virus(es) of each virus are shown on the right. Note that the SDeV phenotypes are shown in gray letters, because there is insufficient information, evidence, or both for the RNA-editing and L-DAg expression. The deltaviruses identified in this study are indicated by the blue circles. Bootstrap values >70 are shown. SDeV: snake deltavirus, RDeV: rodent deltavirus.

### Candidate helper viruses

To gain insights into helper viruses of the novel deltaviruses, we first analyzed the coexisting viruses in the SRA data using BLASTx. Note that we omitted experimental woodchuck hepatitis virus (WHV) infections associated with the mmDeV-positive woodchuck-derived SRA data (SRR2136864 to SRR2136999). We also excluded viruses that infect invertebrates and endogenous retroviruses as well. These analyses reveal that polyomavirus, bornavirus, and circovirus sequences are present in the deltavirus-positive SRA data of passerine birds (Table 3 and Supp Table 4). Further, we detected contigs with 98%–100% identities to human viruses (human mastadenovirus or mammalian rubulavirus 5) (Supp Table 4), although these may represent contamination, index hopping, or both. Among these viruses, only the genome of bornavirus encodes an envelope protein. Note that scDeV and bornavirus-positive SRA data was obtained from pooled samples (SAMN04260514), and we therefore were unable to determine whether scDeV and canary bornavirus 3 infected the same individual.

**Table 3.** Coexisting viruses in deltavirus-positive SRAs

| SRA accession | Host |  | Virus name | Envelope | Deltavirus infection |
| --- | --- | --- | --- | --- | --- |
|  | Species | Common name |  |  |  |
| SRR2545944 | *Taeniopygia guttata* | Zebra finch | Serinus canaria polyomavirus | - | tgDeV |
| SRR5001849 | *Taeniopygia guttata* | Zebra finch | Serinus canaria polyomavirus | - | tgDeV |
| SRR5001850 | *Taeniopygia guttata* | Zebra finch | Serinus canaria polyomavirus | - | tgDeV |
| SRR5001851 | *Taeniopygia guttata* | Zebra finch | Serinus canaria polyomavirus | - | tgDeV |
| SRR2915371 | *Serinus canaria* | Common canary | Canary bornavirus 3 | + | scDeV |
|  |  |  | Canary circovirus | - |  |
| SRR7504989 | *Erythrura gouldiae* | Gouldian finch | Erythrura gouldiae polyomavirus 1 | - | egDeV |

We next cross-referenced the mmDeV reads and the metadata, which also provide insights into the mmDeV helper virus. Among 20 mmDeV-positive SRA data, 18 were obtained from animals experimentally infected with the hepadnavirus WHV, which was experimentally shown to serve as a helper virus for HDV (26, 27). However, the other two SRA data (SRR437934 and SRR437938) were derived from animals negative for antibodies against WHV as well as WHV DNA (24). These observations suggest that mmDeV was transmitted to the two animals without WHV and that WHV therefore was not the helper virus for mmDeV that infected these two individuals.

### Replication of tgDeV and mmDeV in human and woodchuck cell lines

To investigate whether the novel deltavirus sequences are replicable or not, we performed transfection-based assays in cell culture systems. We constructed plasmid expression vectors harboring the minus-strand genome of the tgDeV or mmDeV dimer sequence under the transcriptional control of the CMV promoter (see Materials and Methods). We first determined if the replication initiated by transfecting these plasmids. These plasmids express the minus-strand genome and therefore DAg protein is expressed if the viral genome replicates (19). We transfected the plasmid vectors into Huh7 human hepatic cells and WCH-17 woodchuck hepatic cells and used western blotting (Figs. 6a and b) and immunofluorescence assay (IFA) (Figs. 6c–f) to detect the expression of DAg proteins. Western blotting detected the expected bands (approximately 22 kDa) only in transfected cells (Figs. 6a and b). Note that a single specific band was detected in lysates prepared from each cell type, suggesting that tgDeV and mmDeV expressed only one DAg isoform. Consistent with the above results, specific signals were observed only in the transfected cells in IFA (Figs. 6c and d, red signals). Together, these data suggest that the tgDeV and mmDeV initiated replication from the constructed plasmids in the cell culture system.

**Figure 6.**
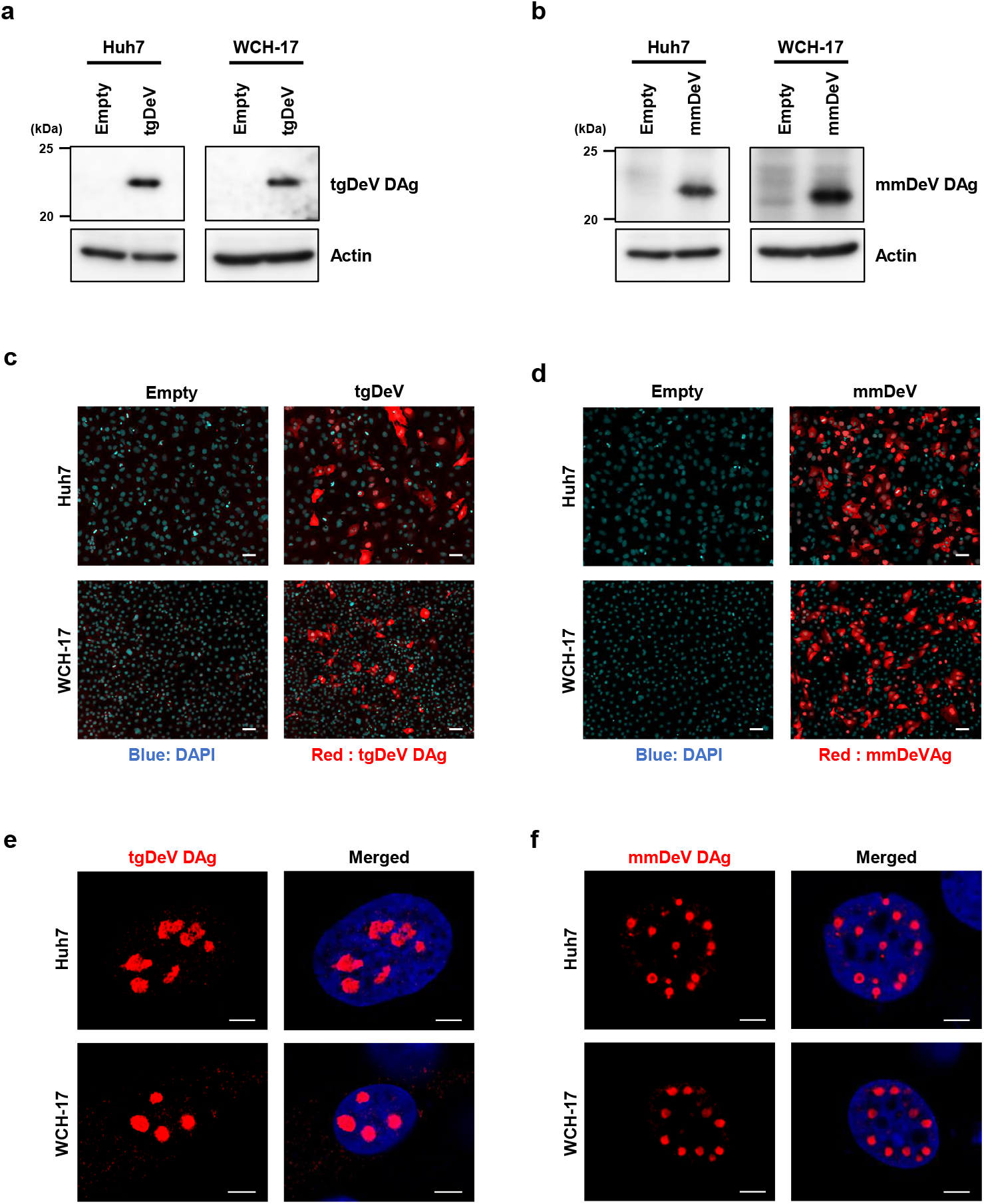
Detection of tgDeV DAg and mmDeV DAg in cells ectopically expressing the tgDeV or mmDeV dimer genome. **(a** and **b)** Western blotting analysis of Huh7 or WCH-17 cells transfected with a tgDeV or mmDeV dimer-sequence expression plasmid. The numbers on the left side of panels indicate the size marker of protein (kDa). **(c–f)** Indirect immunofluorescence analysis of the expression of tgDeV or mmDeV DAg protein. The cells were observed using fluorescent microscopy **(c** and **d)** or a confocal microscopy **(e** and **f)**. Blue; DAPI, Red; tgDeV or mmDeV DAg. Scale bars = 50 μm (c and d) and 5 μm (e and f).

The DAg proteins predominantly localized to the nucleus 2 days post-transfection (Figs. 6e and f). Interestingly, large viral speckles were observed in the nucleus, similar to those detected in cells infected with HDV (28, 29). These results suggest that tgDeV and mmDeV employ a nuclear replication strategy similar to that used by HDV.

### HBV envelope proteins do not contribute to the production of infectious tgDeV and mmDeV

As described in the above section “*Candidate helper viruses*”, there is no evidence of coinfections of hepadnaviruses with tgDeV or mmDeV. However, this does not necessarily mean hepadnaviruses do not serve as helper viruses for the novel deltaviruses. To determine whether tgDeV or mmDeV utilize the HBV envelope proteins (HBs), we transfected the deltavirus expression plasmids together with an HBs expression vector or the cognate empty vector into Huh7 cells. The culture supernatants were incubated with HepG2-NTCP cells, which are susceptible to HBs-dependent HDV infection (30, 31). HDV served as a control to monitor HBs-dependent virus release and subsequent cell entry. Viral RNA was undetected in supernatants of cells that did not express HBs (Fig. 7a). In contrast, cotransfection of the HBs plasmid released large amounts of HDV RNA into the supernatant, consistent with published data (32), whereas tgDeV or mmDeV RNA was undetectable (Fig. 7a).

**Figure 7.**
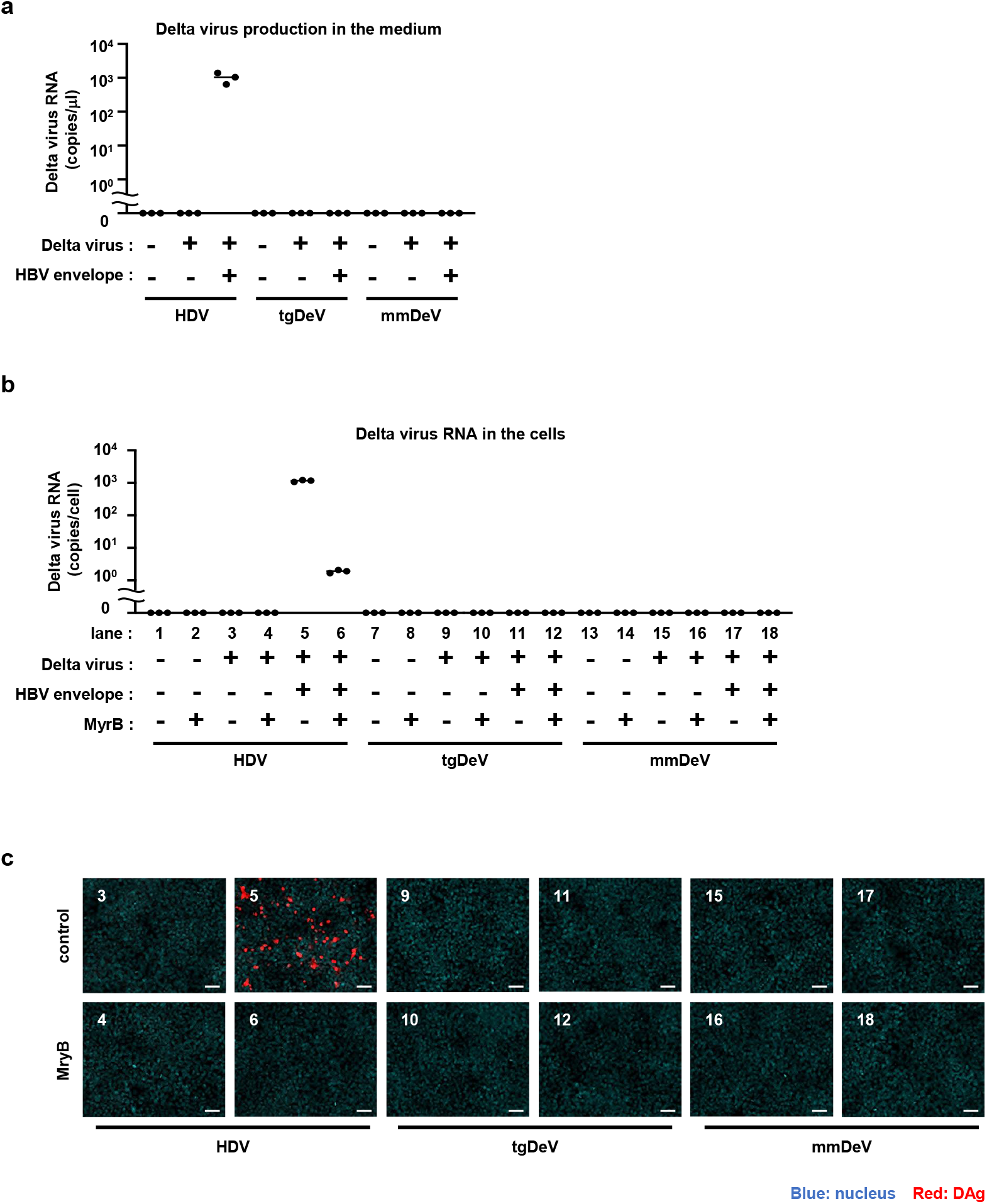
No infectious particle of tgDeV and mmDeV was produced by supplementation of HBV envelop proteins. **(a)** Quantification of deltavirus RNAs in culture supernatants. HDV, tgDeV, or mmDeV expression plasmid was transfected with or without plasmid expressing HBV envelope proteins into Huh7 cells. Viral RNA levels in supernatants were quantified using quantitative RT-PCR (n = 3). **(b** and **c)** HepG2-NTCP cells were incubated with the culture supernatants of the transfectants for 24 h in the presence or absence of 500 nM Myrcludex B (MyrB), an inhibitor of HBV envelope-dependent viral entry. The cells were cultured for an additional 6 days, and viral RNA levels and protein exprssion were analyzed using quantitative RT-PCR (n = 3) **(b)** and IFA **(c)**, respectively. The numbers in (**c**) correspond to those of (**b**). Blue, DAPI; Red, HDV; tgDeV, or mmDeV DAg. Scale bar = 50 μm.

We next measured the amounts of viral RNA and detected DAg protein in HepG2-NTCP cells 7 days after incubation with the supernatants. HDV RNA and DAg protein were highly expressed in the infected cells (Fig. 7b and c). HDV infection was inhibited by the preS1 peptide (Myrcludex B), which was shown to inhibit HBs-dependent HDV infection (33). These indicate that HDV infection is indeed mediated by the HBs. On the other hand, tgDeV and mmDeV RNA or DAg protein was undetectable, suggesting that HBs do not contribute to the production of infectious tgDeV or mmDeV.

## Discussion

Important aspects of the evolution of deltaviruses are unknown, such as the origin of HDV and the coevolution of deltaviruses and their helper viruses, mainly because few deltaviruses are known, and they are highly genetically divergent (15–18). Therefore, the resulting large gaps in the deltavirus phylogenetic tree create a formidable obstacle to understanding deltavirus evolution. Here we identified five complete genomes of novel deltaviruses from birds and mammals (Fig. 1 and Supp Fig. 5), which partially fill these phylogenetic gaps (Fig. 5b). Moreover, our present findings reveal that the evolution of deltaviruses is much more complicated than previously thought. For example, one hypothesis states that mammalian deltaviruses codiverged with their mammalian hosts (18). However, our phylogenetic analysis shows that the tree topology of mammalian deltaviruses is incongruent with their hosts’. For example, ovDeV, which we detected in deer, is most closely related to human HDV (Fig. 5b). Further, the distantly related mmDeV and rodent DeV (detected in *Proechimys semispinosus* (18)) were detected in rodent species. These data suggest that deltaviruses were transmitted among mammalian species and did not always codiverge with their hosts. Moreover, we discovered recent interfamily transmission of passerine deltaviruses (tgDeV and its relatives) (Fig. 4e). Therefore, avian and mammalian deltaviruses may have, at least partially, evolved by interspecies transmission.

Our present phylogenetic analysis also gives insights into the origin of human HDVs. As described above, ovDeV and mmDeV are close relatives of human HDVs, suggesting that HDVs arose from other mammalian deltaviruses. Recent studies on the phylogeny of bat deltaviruses support our findings and conclusions (34, 35) (Supp Fig. 7). However, we were unable to exclude the possibility of infection of animal lineages apart from mammals with unknown deltaviruses phylogenetically located between those viral lineages. Further investigations are required for a better understanding of the deltavirus evolution.

There is a paucity of knowledge about helper viruses for non-HDV deltaviruses, other than the snake deltavirus (19), although evidence indicates that hepadnaviruses may not serve as helper viruses for the novel deltaviruses discovered here, as suggested for other non-HDV deltaviruses (15–18). Here we only detected bornavirus, circovirus, and polyomavirus, but not hepadnavirus sequences in association with deltavirus-positive SRA data (Table 3). Further, mmDeV was detected in two woodchuck individuals that were demonstrated to be negative for WHV (Table 2 and Supp Table 3). Moreover, we found that HBs did not contribute to the formation of infectious tgDeV and mmDeV in cell culture experiments. These observations suggest that hepadnaviruses do not serve as helper viruses for the novel non-HDV deltaviruses detected here. Further, we were unable to demonstrate that the deltaviruses identified here express proteins similar to the L-HDAg protein (Figs. 6a and b), which is expressed via RNA-editing and is essential for HDV to interact with HBs (2, 37). Although RNA-editing may alter the stop codon of ovDeV DAg, this does not lead to the expression of a large isoform of DAg (L-DAg) protein containing a farnesylation site (Figs. 2b and c). The lack of L-DAg expression was also observed in rodent deltaviruses (18). Therefore, L-DAg expression phenotype may have been acquired after the divergence of ovDeV and the HDV lineage (Fig. 5b).

Among the coexisting viruses, only bornavirus produces an envelope glycoprotein (G protein), which might be used by non-HDV deltaviruses to produce infectious virions. Indeed, snake deltavirus utilizes the envelope proteins of reptarenaviruses and hartmaniviruses to produce infectious particles (19). Furthre, HDV forms infectious virions using envelope proteins *in vitro* of RNA viruses such as vesiculovirus and hepacivirus (38). Therefore, the bornavirus G protein might envelop non-HDV deltaviruses.

In contrast, the coexisting viruses, circoviruses and polyomaviruses, are nonenveloped. Therefore, it is unlikely that these viruses can serve as helper viruses for the deltaviruses. However, we cannot exclude the possibility that these viral capsid proteins might contribute to the transmissibility of deltaviruses through unknown mechanisms. Additionally, virus-derived sequences in host genomes, called endogenous viral elements (EVEs), might mediate the formation of infectious particles. Here detected the expression of retroviral EVEs in certain deltavirus-positive SRA data (data not shown). Although HDVs do not use retroviral envelope proteins (38), non-HDV deltaviruses might utilize strategies distinct from those employed by HDVs. Alternatively, non-HDV deltaviruses may not require helper viruses and utilize extracellular vesicles for transmission. Further biological experiments, together with molecular surveillance, are therefore required to understand the satellite-helper relationships of deltaviruses.

Here we show that the sequences of tgDeV and tgDeV-like viruses, such as pvDeV and lsDeV, are relatively closely related to known vertebrate deltaviruses (Fig. 5b). Although a previous study found a deltavirus from duck, this duck-associated virus was detected in oropharyngeal/cloacal swabs and is distantly related to vertebrate deltaviruses, suggesting the possibility of its dietary origin (15, 18). This may be true for scDeV studied here. scDeV was detected in skin (Table 1). Although scDeV was excluded from our phylogenetic analysis, the amino acid identities between the DAg protein of scDeV with other vertebrate deltaviruses range from 32.7%–39.5% (Fig. 5a). Therefore, scDeV may be derived from contaminants, which should be addressed in the future. In contrast, tgDeV and tgDeV-like viruses were detected in tissues such as the spleen and muscles (Table 2), suggesting that tgDeV and tgDeV-like viruses are authentic avian deltaviruses.

Here we show that certain novel deltaviruses are transmitted among animal populations (Table 2 and Supp Table 3). Note that few reads were mapped to the virus genomes in some SRA data, which may be attributed to index hopping (39–43) from SRA data containing numerous deltavirus-derived reads. Therefore, these data should be interpreted with caution. Nevertheless, our conclusions are not affected, because they are supported by robust data (Table 2). For data in which index hopping has possibly occurred, further analyses are needed to confirm deltavirus infections.

Our present analysis provides virological insights into important characteristics of deltavirus infections, such as tissue and host tropism. For example, infections with tgDeV (and tgDeV-like viruses), mmDeV, and ovDeV were not limited to the liver and were detected in at least two different tissues (Table 2). These observations are consistent with those of previous studies that non-HDV deltaviruses in multiple organs and blood and that they replicate in numerous cell types (18, 19). Therefore non-HDV deltaviruses may infect diverse tissues and cause systemic infection, viremia, or both. Furthre, tgDeV and mmDeV replicated in human and woodchuck cells (Fig. 6), which is consistent with the ability of the snake deltavirus to replicate in mammalian cells (19). These observations suggest that the host range of deltaviruses is broad and that the helper viruses of non-HDV deltaviruses may be the determinants of host range.

Our analyses further suggest that tgDeV and mmDeV are sensitive to host immune responses. We cross-referenced tgDeV reads and metadata and made an intriguing finding that may contribute to the virus-host interaction. BioProject PRJNA297576 contains 12 RNA-seq data for six zebra finches (44). Interestingly, the tgDeV reads were almost exclusively detected in birds treated with testosterone vs the controls (Supp Fig. 8a and Supp Table 5). Therefore, the immunosuppressive effects of testosterone (45) may increase the transcription or replication of tgDeV, or both, to enable detection using RNA-seq. Furthre, when we cross-referenced the mmDeV reads and the metadata, we found that 18 of 20 mmDeV-positive SRA data derived from five individuals were acquired through an experiment lasting 27 weeks (PRJNA291589) (25). Among 18 SRA data, those of one individual (ID 1008) provide insights into mmDeV infection as follows: At first (–3 weeks), mapped reads were not detected, although the proportion of mapped reads were highest at one week and then drastically decreased (Supp Fig. 8b and Supp Table 6). Interestingly, a previous study suggested that the host’s immune response can clear rodent deltaviruses (18). Our present observations together with this previous finding, suggest that the host immune response suppresses and then clear deltavirus infections. This may explain the low prevalence of RT-PCR-positive samples of woodchucks and passerine birds as described in the Results (sections “*Circulation of tgDeV and tgDeV-like viruses among passerine birds*” and “*Circulation of mmDeV in woodchucks*”). Note that latent or low levels of persistent deltavirus infections may occur. Indeed, snake deltavirus establishes a persistent infection in a cell culture system (19). Therefore, deltaviruses might persistently infect host cells with a low level of virus replication, and some stimulations, such as immunosuppression, may trigger robust virus replication. Further studies are therefore required to understand the interactions between deltaviruses and their hosts.

Together, our present data contribute to a deeper understanding of the evolution of deltaviruses and suggest the presence of undiscovered deltaviruses that infect diverse animal species. Further investigations will provide further insights into deltavirus evolution.

## Materials and methods

### Detection of deltaviruses from publicly available transcriptome data

Paired-end, RNA-seq data from birds and mammals were downloaded from NCBI SRA (46). The SRA accession numbers used in this study are listed in Supplementary material. The downloaded SRA files were dumped using pfastq-dump (DOI: 10.5281/zenodo.2590842; https://github.com/inutano/pfastq-dump), and then preprocessed using fastp 0.20.0 (47). If genome data of either the same species or the same genus were available, the preprocessed reads were mapped to the corresponding genome sequences (the genome information is available upon request) by HISAT2 2.1.0 (48), and then unmapped paired-end reads were extracted using SAMtools 1.9 (49) and Picard 2.20.4 (http://broadinstitute.github.io/picard/). The extracted unmapped reads were used for *de novo* assembly. If genome data were unavailable, the preprocessed reads were directly used for *de novo* assembly. *De novo* assembly was conducted using SPAdes (50) and/or metaSPAdes (51) 3.13.0 with k-mer of 21, 33, 55, 77, and 99. The resultant contigs were clustered by cd-hit-est 4.8.1 (52, 53) with a threshold of 0.95. Finally, the clustered contigs ≥ 500 nt were extracted by SeqKit 0.9.0 (54), and they were used for the downstream analyses.

Two-step sequence similarity searches were performed to identify RNA virus-like sequences. First, BLASTx searches were performed against a custom database for RNA viruses using the obtained contigs as queries employing BLAST+ 2.9.0 (55) with the following options: -word_size 2, -evalue 1e^−3^, max_target_seqs 1. The custom database of RNA viruses consisted of clustered sequences (by cd-hit 4.8.1 with a threshold of 0.98) from viruses of the realm *Riboviria* in the NCBI GenBank (the sequences were downloaded on June 2, 2019) (46). Next, the query sequences with viral hits were subjected to second BLASTx analyses, which were performed against the NCBI nr database. Finally, the second blast hits with the best hit against deltaviruses were regarded as deltavirus-like agents, and they were used for detailed analyses.

### Confirmation of circularities of deltavirus contigs

Self dot-plot analyses of linear deltavirus contigs were conducted using the YASS online web server (29). Based on the analysis, the contigs were manually circularized using Geneious 11.1.5 (https://www.geneious.com). Further confirmation of the circularities of deltavirus contigs was obtained by mapping short reads to circular deltavirus contigs using Geneious software as follows. The reads used for the *de novo* assembly were first imported to Geneious, after which they were mapped to the circular contigs using the Geneious mapper. The mapped reads across the circularized boundaries were confirmed manually.

### Detection of possible RNA-editing sites at stop codons of DAg genes

We used the mapped reads obtained by the analyses described above to detect possible RNA-editing at the stop codons of DAg genes of deltaviruses. We analyzed the nucleotide variations (presence of variations, variant nucleotide(s), and variant frequency) of mapped reads at each of the stop codons of newly identified deltaviruses using the “Find Variation/SNPs” function in Geneious. We used a custom Python script to visualize base quality scores of the NGS reads mapped at the second nucleotide of the stop codon genome. The codes are available at following URL: https://github.com/shohei-kojima/iwamoto_et_al_2020.

### Sequence characterization

DAg ORFs were detected by the “Find ORFs” function in Geneious with a threshold of 500 nucleotides. Poly(A) signals were manually detected. Putative ribozyme sequences were identified using nucleotide sequence alignment with other deltaviruses. Ribozyme structures were first inferred using the TT2NE webserver (56), and the obtained data were then visualized using the PsudoViewer3 web server (57). We used the visualized data as guides to draw ribozyme structures.

The self-complementarities of deltavirus-like contigs were analyzed using the Mfold web server (58). Coiled-coil domains and nuclear export signals (NLSs) were predicted using DeepCoil (59) and NLS mapper (http://nls-mapper.iab.keio.ac.jp/cgi-bin/NLS_Mapper_form.cgi) web servers, respectively.

### Short read mapping for detection of deltavirus infection

To detect deltavirus-derived reads in publicly available RNA-seq data, short reads were mapped to deltavirus genomes and then the numbers of mapped reads were counted as follows. SRA files were downloaded from NCBI, dumped, and preprocessed following the procedure described above. The preprocessed reads were then mapped to linearized deltavirus contigs by HISAT2 with the default setting. SAM tools were used to extract the mapped BAM files from the resultant BAM files, and the mapped read numbers were counted using BamTools 2.5.1 (60).

### Recovery of a deltavirus genome from RNA-seq data of Pardaliparus venustulus

Mapped reads obtained from SRR7244693, SRR7244695, SRR7244696, SRR7244697, and SRR7244698 in the above analysis (section *Short read mapping for detection of deltavirus infection*) were extracted by Geneious. All the extracted reads were co-assembled using Geneious Assembler with the circular contig assembly function. The obtained circular contigs were characterized as described previously.

### Animals and samples

Zebra finches (n = 30) and Bengalese finches (n = 5) were obtained from breeding colonies at Wada lab, Hokkaido University. The founder birds were originally obtained from local breeders in Japan. Five to twelve birds were kept together in cages in an aviary and were exposed to a 13:11 light-dark cycle. Blood samples were collected from the wing vein using 30 G × 8 mm syringe needles (Becton Dickinson; Franklin Lakes, NJ, USA). Each blood sample was diluted 1.5 times with PBS, frozen immediately on dry ice after collection, and maintained at −80°C until further requirement. These experiments were conducted under the guidelines and with the approval of the Committee on Animal Experiments of Hokkaido University. These guidelines are based on the national regulations for animal welfare in Japan (Law for the Humane Treatment and Management of Animals with partial amendment No.105, 2011).

Woodchucks (*Marmota monax*) were purchased from Northeastern Wildlife (Harrison, ID, USA) and kept at the Laboratory Animal Center, National Taiwan University College of Medicine. At three days of age, the animal supplier infected captive-born woodchucks with WHV from the same infectious pool. Wild-caught woodchucks were infected naturally and live trapped. Serum samples were collected from the woodchucks periodically via the femoral vein by means of venipuncture. Liver tissues of woodchucks were obtained at autopsy, snap-frozen in liquid nitrogen, and stored at −80°C until RNA extraction. This study used liver tissues from 10 wild-caught and 33 captive-born woodchucks and serum samples from 33 wild-caught and five captive-born woodchucks. In this study, all the experimental procedures involving woodchucks were performed under protocols approved by the Institutional Animal Care and Use Committee of National Taiwan University College of Medicine.

### Real-time and Endpoint RT-PCR detection of deltaviruses from animal specimens

Total RNAs were isolated from the whole blood samples from zebra finches and serum samples from woodchucks using Quick RNA Viral Kit (Zymo Research; Irvine, CA, USA). The obtained RNA samples were stored at −80°C until further requirement. Total RNAs were also extracted from 50 mg of the woodchuck liver tissues using either Trizol (Thermo Fisher Scientific; Waltham, MA, USA) or ToTALLY RNA kit (Thermo Fisher Scientific; Waltham, MA, USA) according to the manufacturers’ instructions.

The obtained RNA was reverse-transcribed into cDNA using ReverTra Ace qPCR RT Master Mix (TOYOBO; Osaka, Japan), and these were used as templates for real-time PCR analyses. Real-time PCR was performed with KOD SYBR qPCR Mix (TOYOBO) and primers (Supp Table 7) using the CFX Connect Real-Time PCR Detection System (Bio-Rad; Hercules, CA, USA) according to the manufacturer’s instructions. The real-time PCR systems for mmDeV and tgDeV were validated using pcDNA3-mmDeV(-) and pcDNA3-tgDeV(-) monomer, respectively, as controls.

End-point RT-PCR was also performed to confirm deltavirus infections. PCR was performed with Phusion Hot Start II DNA Polymerase (Thermo Fisher Scientific) using the above-described cDNAs and primers listed in Supp Table 7. The PCR products were analyzed by agarose gel electrophoresis. The obtained PCR products were purified and sequenced by Sangar sequencing in FASMAC (Atsugi, Japan).

### Determination of a full genome sequence of deltavirus in passerine birds

To determine the full genome sequence of detected deltaviruses, the cDNA obtained in the section *“Realtime and Endpoint RT-PCR detection of deltaviruses from animal specimens*” was amplified using illustra GenomiPhi V2 Kit (GE healthcare; Chicago, IL, USA). The amplified DNA was then purified with innuPREP PCRpure Kit (Analytik Jena: Jena, Germany). PCR was performed with Phusion Hot Start II DNA Polymerase using the primers listed in Supp Table 7. The PCR products were analyzed using agarose gel electrophoreses. When single bands were observed, the amplicon was purified with innuPREP PCRpure Kit. When several bands were detected, bands of the expected sizes were extracted and purified using Zymoclean Gel DNA Recovery Kit (Zymo Research). The purified amplicons were sequenced in FASMAC (Atsugi, Japan).

### Phylogenetic analysis

Deduced amino acid sequences of DAg proteins were used to infer the phylogenetic relationship between deltaviruses. Multiple alignment was performed by MAFFT 7.427 using the E-INS-i algorithms (61), and ambiguously aligned regions were then removed using trimAl 1.2rev59 with the --strict option (62). The phylogenetic relationship was inferred by the maximum likelihood method using RAxML Next Generation v. 0.9.0 (63). The LG+G model, which showed the lowest BIC by prottest3 3.4.2 (64), was used. The reliability of the tree was assessed by 1,000 bootstrap resampling using the transfer bootstrap expectation method (65). The alignment file is available in Supporting materials.

### Detection of co-infected viruses

To identify co-infected viruses in deltavirus-positive SRAs, a three-step BLASTx search was performed. First, BLASTx searches were performed against a custom database, including RefSeq protein sequences from viruses using the assembled contigs (see the subsection *Detection of deltaviruses from publicly available transcriptome data*) as queries. The custom database was prepared as follows. Virus-derived protein sequences in the RefSeq protein database (46) were downloaded on July 17, 2020, and were clustered by cd-hit 4.8.1 (threshold = 0.9). Then, sequences of more than 100 amino acid residues were extracted using SeqKit 0.10.1 and these were used as a BLAST database. The first BLAST hits were extracted, which were used for the second BLASTx analysis. The second BLASTx analysis was performed against the NCBI RefSeq protein database. The BLAST hits with the best hit to viral sequences were extracted and used for the final BLASTx searches. The final BLASTx searches were performed against the NCBI nr database. The BLAST hits with the best hit to viral sequences were extracted and analyzed manually.

### Cell culture

HepG2-NTCP cells were cultured with Dulbecco’s modified Eagle’s medium (DMEM)/F-12 + GlutaMax (Thermo Fisher Scientific) supplemented with 10 mM HEPES (Sigma Aldrich; St. Louis, MO, USA), 100 unit/ml penicillin (Meiji; Tokyo, Japan), 100 mg/ml streptomycin (Meiji), 10% FBS (Sigma Aldrich), 5 μg/ml insulin (Wako; Tokyo, Japan) and 400 g/ml G418 (Nacalai tesque). Huh7 and WCH-17 cells were maintained in DMEM (Wako) containing 10% FBS (Sigma Aldrich), 100 unit/ml penicillin (Meiji), 100 mg/ml streptomycin (Meiji), 100 mM nonessential amino acids (Thermo Fisher Scientific), 1 mM sodium pyruvate (Sigma Aldrich), and 10 mM HEPES (Sigma Aldrich).

### Antibody production

The peptides corresponding to 65 to 78 aa (DSSSPRKRKRGEGG) of tgDeV DAg and 174 to 187 aa (ESPYSRRGEGLDIR) of mmDeV DAg conjugated with cysteine at N terminus were synthesized. Each of the peptides was injected into mice, and antisera were obtained from the mice at 42 days after the peptide injections. Each of the antisera was affinity-purified using the corresponding peptide. The whole procedure was performed in SCRUM (Tokyo, Japan).

### Rescue of mmDeV and tgDeV

The DNA of negative-strand genomes of mmDeV and tgDeV was synthesized in GenScript Japan (Tokyo, Japan). The synthesized DNAs were inserted into the KpnI -XbaI site of the pcDNA3 vector, designated as pcDNA3-mmDeV(-) monomer and pcDNA-tgDeV (-) monomer. In addition, tandem sequences of mmDeV and tgDeV genome were inserted into the pcDNA3 vector, which were named pcDNA3-mmDeV(-) dimer and pcDNA-tgDeV (-) dimer, respectively. To rescue these viruses, pcDNA3-mmDeV(-) dimer or pcDNA-tgDeV (-) dimer was transfected into Huh7 and WCH-17 cells using Lipofectamine 3000 and Lipofectamine 2000 (Thermo Fisher Scientific), respectively, according to the manufacturer’s instructions. The transfected cells were cultured for 48 h and were used for western blotting, IFA to verify DAg protein expression.

### Western blotting

Cells were lysed with SDS sample buffer [100 mM Tris-HCl (pH 6.8) (Sigma Aldrich), 4% SDS (Nippon gene; Tokyo, Japan), 20% glycerol (Nacalai tesque), 10% 2-mercaptoethanol (Wako)]. The cell lysates were subjected to SDS-PAGE and transferred onto polyvinylidene difluoride membranes (Merck Millipore; Darmstadt, Germany). After blocking the membranes with 5% skim milk (Morinaga; Tokyo, Japan), they were reacted with anti-tgDeV DAg, anti-mmDeV DAg, or anti-actin (Sigma Aldrich) antibodies as primary antibodies, followed by reaction with horseradish peroxidase (HRP)–conjugated secondary antibodies (Cell Signaling Technology; Danvers, MA, USA).

### Indirect immunofluorescence assay (IFA)

The cells were fixed in 4% paraformaldehyde (Wako) and then permeabilized using 0.3% Triton X-100 (MP Biomedicals; Santa Ana, CA, USA). After blocking the cells by incubation in PBS containing 1% bovine serum albumin (BSA) (KAC; Kyoto, Japan), they were treated with the primary antibodies against HDAg, tgDeV DAg, or mmDeV DAg and then incubated with Alexa555-conjugated secondary antibody (Thermo Fisher Scientific), together with DAPI (Nacalai tesque). To detect deltavirus-positive cells, the fluorescence signal was observed using fluorescence microscopy, BZ-X710 (KEYENCE; Osaka, Japan). High magnification examination of the subcellular localization was performed using confocal microscopy, LSM900 (ZEISS; Oberkochen, Germany).

### Deltavirus preparation and infection assay

HDV was produced from the culture supernatants of Huh7 cells transfected with HDV (pSVLD3) and HBs (pT7HB2.7) expressing plasmid, as described previously (32, 66). tgDeV and mmDeV were also subjected to the same assay. The supernatants of transfected cells were collected at 6, 9, and 12 days post-transfection, and they were then filtrated and concentrated using 0.45-μm filters and Amicon Ultra (Merck Millipore), according to the manufacturer’s instructions. The concentrated supernatants were inoculated into HepG2-NTCP cells with 5% PEG8000 (Sigma Aldrich) for 24 h followed by washing to remove free viruses. The inoculated cells were cultured for 6 days and used for the downstream analyses.

## Acknowledgments

HDV and HBs expression plasmids were kindly provided by Dr. John Taylor (the Fox Chase Cancer) and Dr. Camille Sureau (Institute National de la Transfusion Sanguine). We thank Dr. Keiko Takemoto (Kyoto University) for her kind help with the setting of computer resources. The super-computing resources were provided by Human Genome Center, the Institute of Medical Science, the University of Tokyo, and the NIG supercomputer at ROIS National Institute of Genetics. All the silhouette images except for woodchuck were downloaded from silhouetteAC (http://www.silhouette-ac.com/).

This study was supported by the Hakubi project at Kyoto University (MH); Grant-in-Aid for Scientific Research on Innovative Areas from the Ministry of Education, Culture, Science, Sports, and Technology (MEXT) of Japan, Grant Numbers JP19H04839 (SI), JP16H06429 (KT), JP16K21723 (KT), JP16H0643 (KT), JP17H05821 (MH), and JP19H04833 (MH); the Japan Society for the Promotion of Science KAKENHI, Grant Numbers JP19K16672 (MI), JP20J00868 (MI), and JP20H03499 (KW); AMED Grant Numbers JP20jm0210068j0002 (KW) and JP20fk0310114j0004 (KW).

## Author contributions

MH conceived the study. MI, KW, and MH designed the study. MI conducted cell culture experiments. MH, JK, SK performed *in silico* analyses. YS, YTL, HLW, KW, and MH prepared and analyzed animal specimens. All the authors analyzed and discussed the data. MI and MH wrote the manuscript.

