## Supplementary Figure for "Identification of novel avian and mammalian deltaviruses provides new insights into deltavirus evolution"

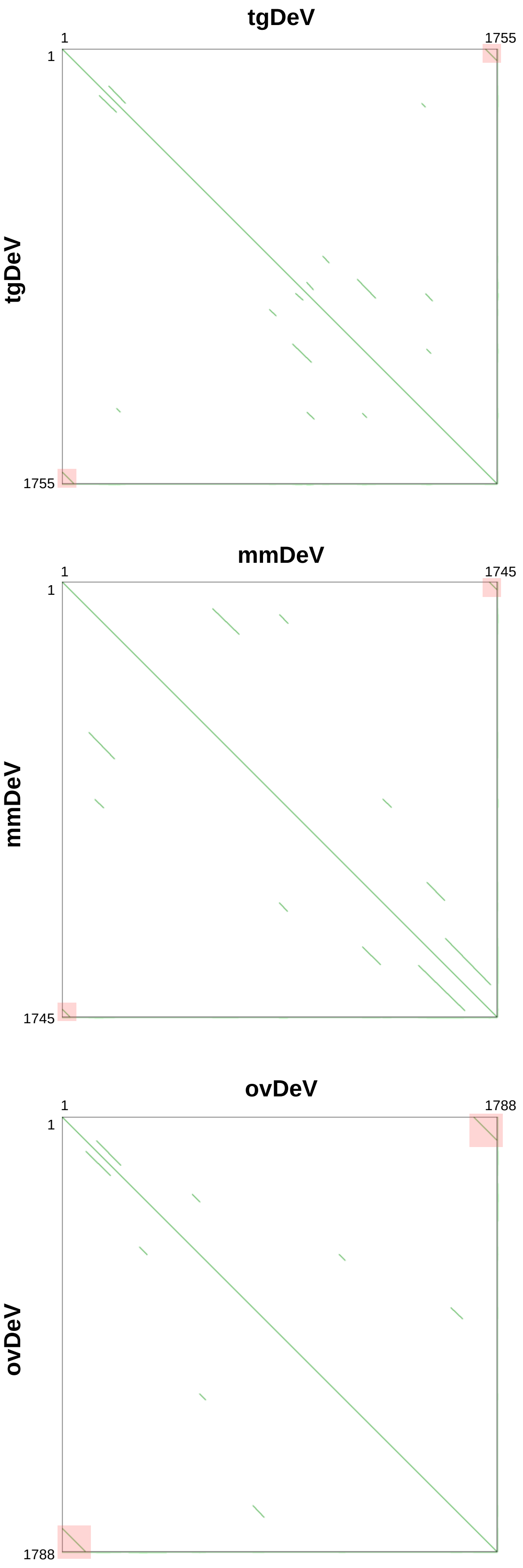


**Supplementary Figure 1. Self-dotplot analyses of deltavirus-like contigs.**
Deltavirus-like contigs were analyzed using the YASS web server. The virus names are shown above and on the left-side of each dot plot. Numbers indicate nucleotide positions. Green lines indicate aligned sequences. Contig ends with identical sequences are highlighted by light-pink boxes.

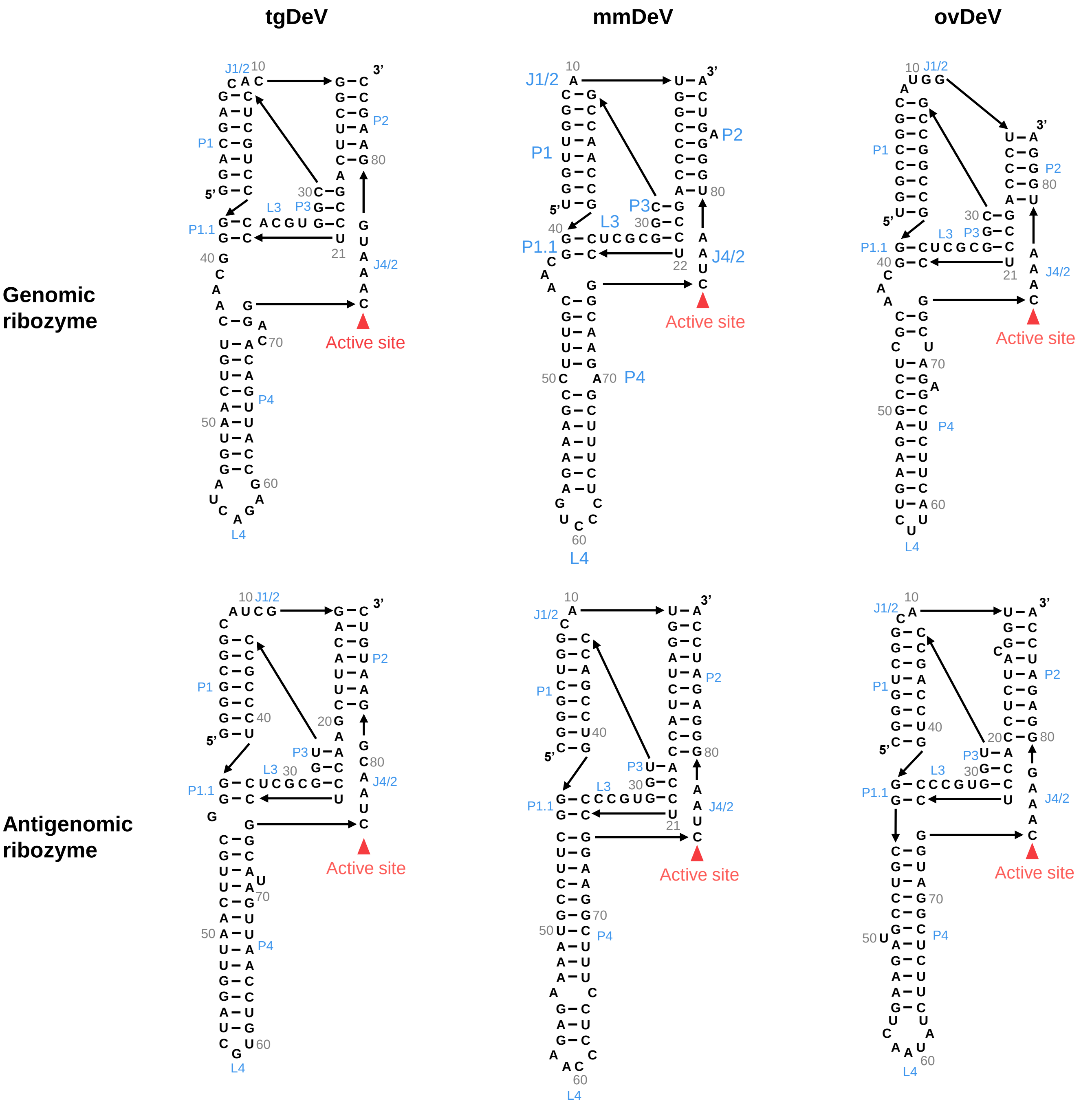


**Supplementary Figure 2. Predicted structures of ribozymes encoded by deltavirus genomes and antigenomes.**

Predicted genomic and antigenomic ribozyme structures of tgDeV, mmDeV, and ovDeV. Gray numbers indicate nucleotide positions of each predicted ribozyme. The names of secondary structural elements of ribozymes are blue. Catalytic sites are indicated by the pink triangles.

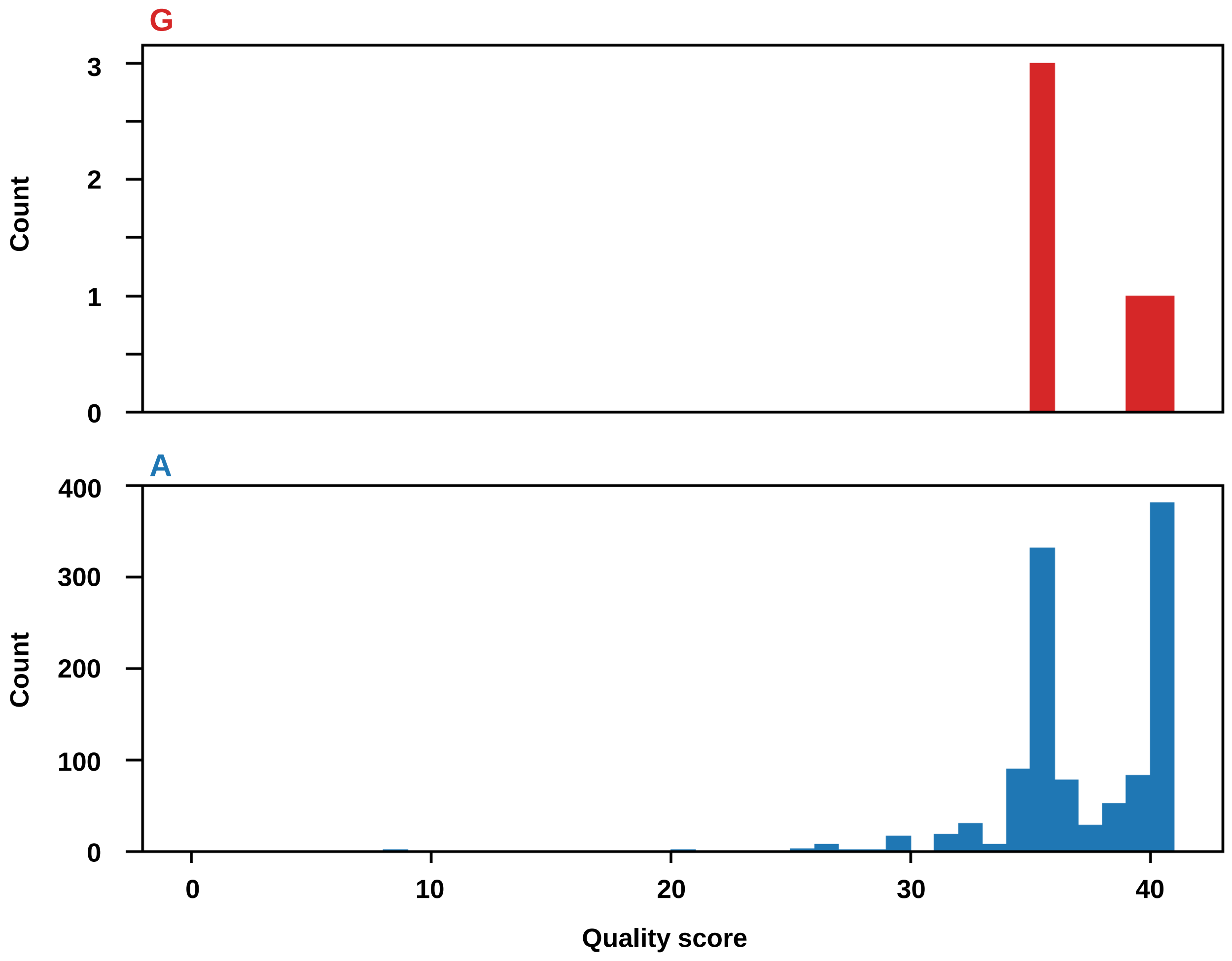


**Supplementary Figure 3. Quality scores of a potential RNA-editing site in ovDeV.** Histograms of the quality scores at the potential RNA-editing site (the second nucleotide position of the stop codon) in the ovDeV-DAg gene of mapped reads. Red and blue histograms indicate the quality scores of G and A nucleotides at this position.

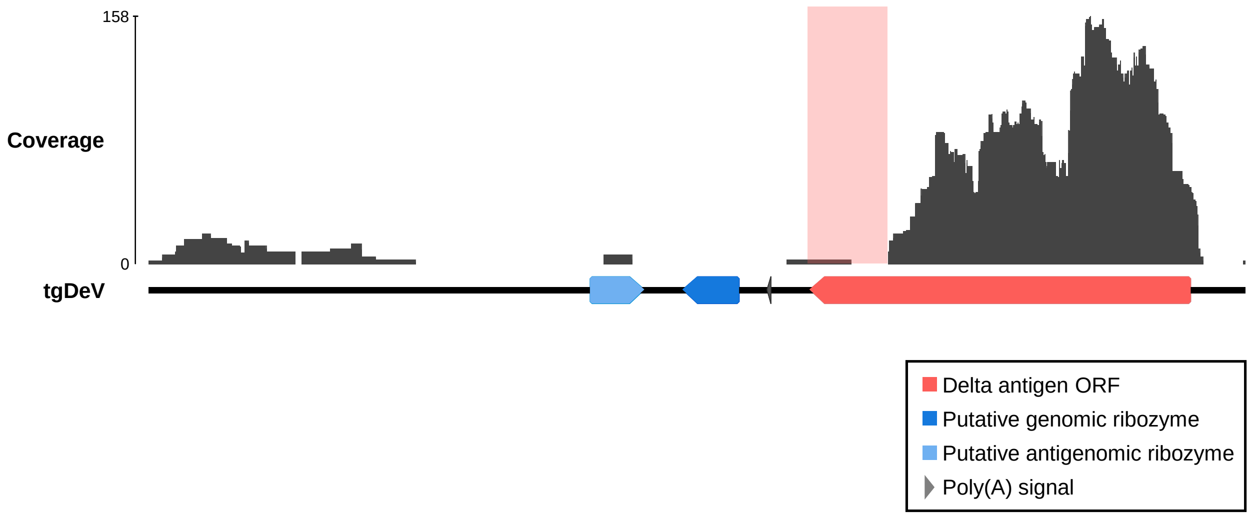


**Supplementary Figure 4. Mapping coverage of Taeniopygia guttata-associated deltavirus.** Mapped reads of tgDeV. Short reads in SRR2545943 that were not mapped to the genome of Taeniopygia guttata were mapped to the tgDeV genome. The line, arrow pentagons, and arrowheads indicate the viral genome, ribozymes, the Dag-ORF, and the poly(A) signal, respectively. The light-pink box shows the low-coverage region of the ORF.

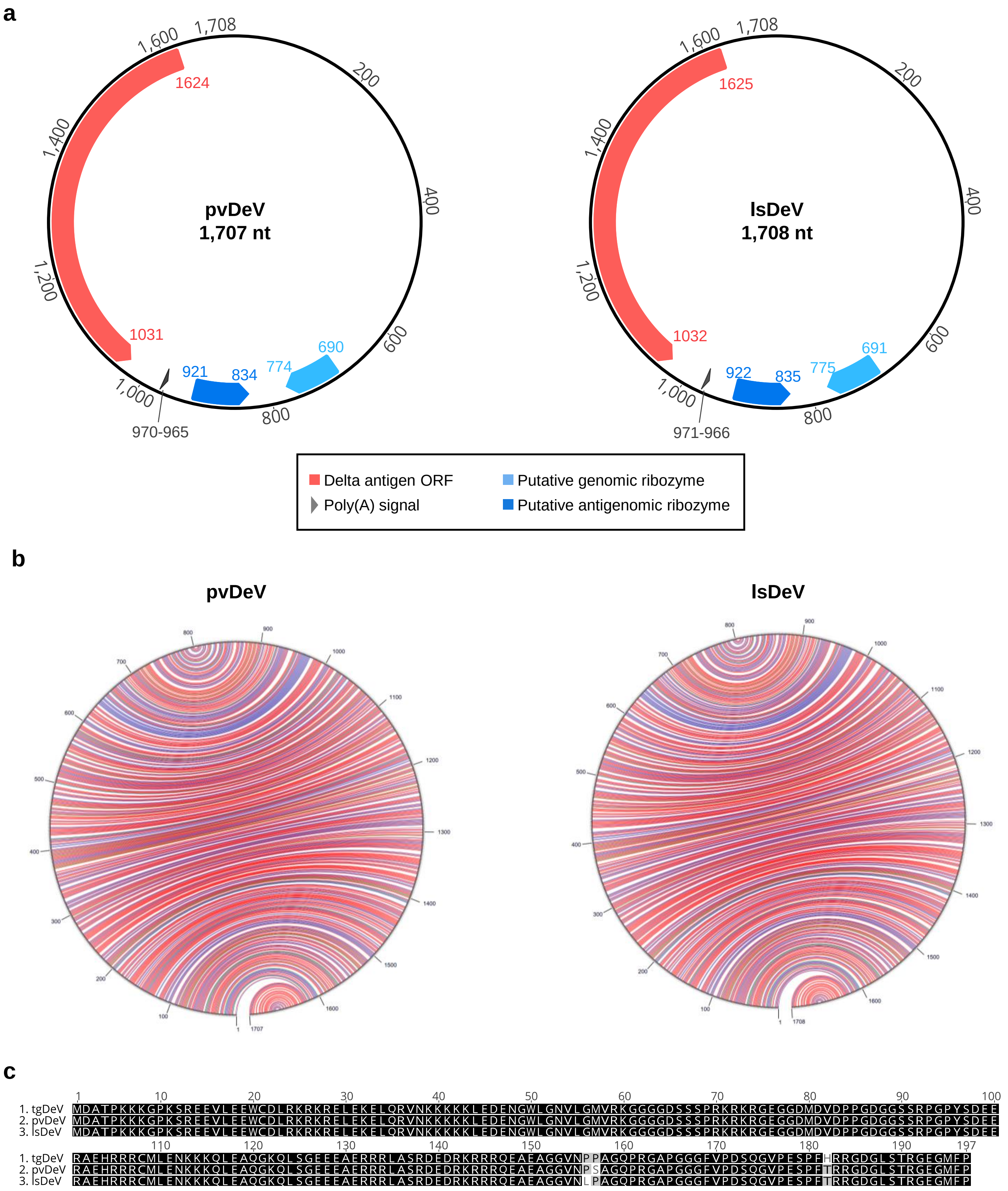


**Supplementary Figure 5. Characterization of pvDeV and lsDeV.**
**(a)** Genome organizations of pvDeV and lsDeV. Annotations are shown by colored pentagons. **(b)** Self-complementarity of the pvDeV and lsDeV genomes. Circular structure plot of pvDeV (mfold web server). Red, blue, and green arcs indicate G-C, A-U, and G-U pairs, respectively. **(c)** Pairwise amino acid sequence alignment of DAg proteins of tgDeV, pvDeV, and lsDeV. The black boxes with white letters indicate identical amino acid resides. The numbers on the alignment indicate the positions.

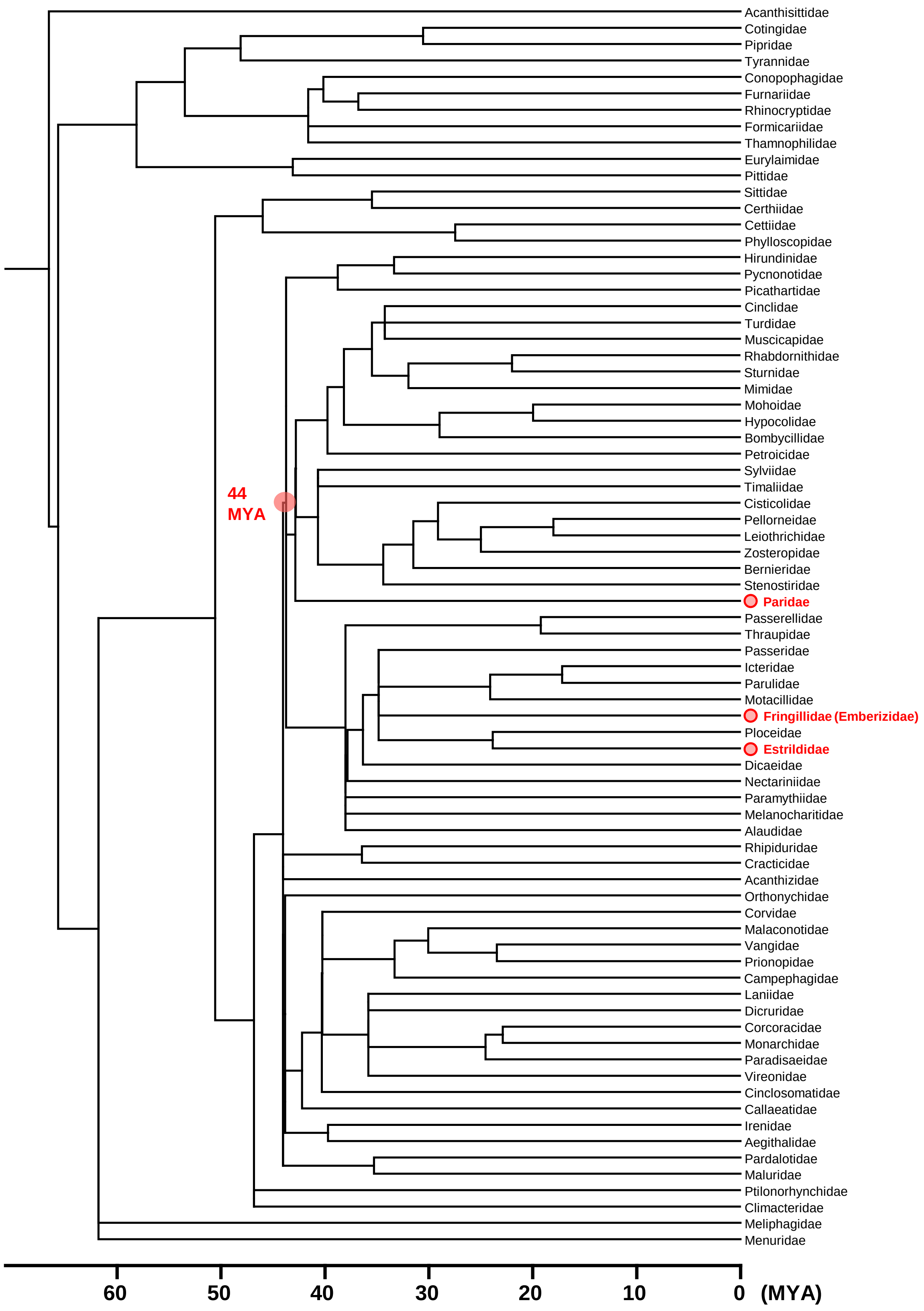


**Supplementary Figure 6. Divergence times of the order Passeriformes.**
A TimeTree phylogenetic tree of the genera in the order Passeriformes is shown. The time scale is shown under the phylogenetic tree. The families in which deltaviruses were detected are indicated by pink circles. The divergence time was approximately to 44 million years ago (MYA) (pink circle). Note that the family Emberizidae is considered a subfamily of Emberizinae of the family Fringillidae in TimeTree.

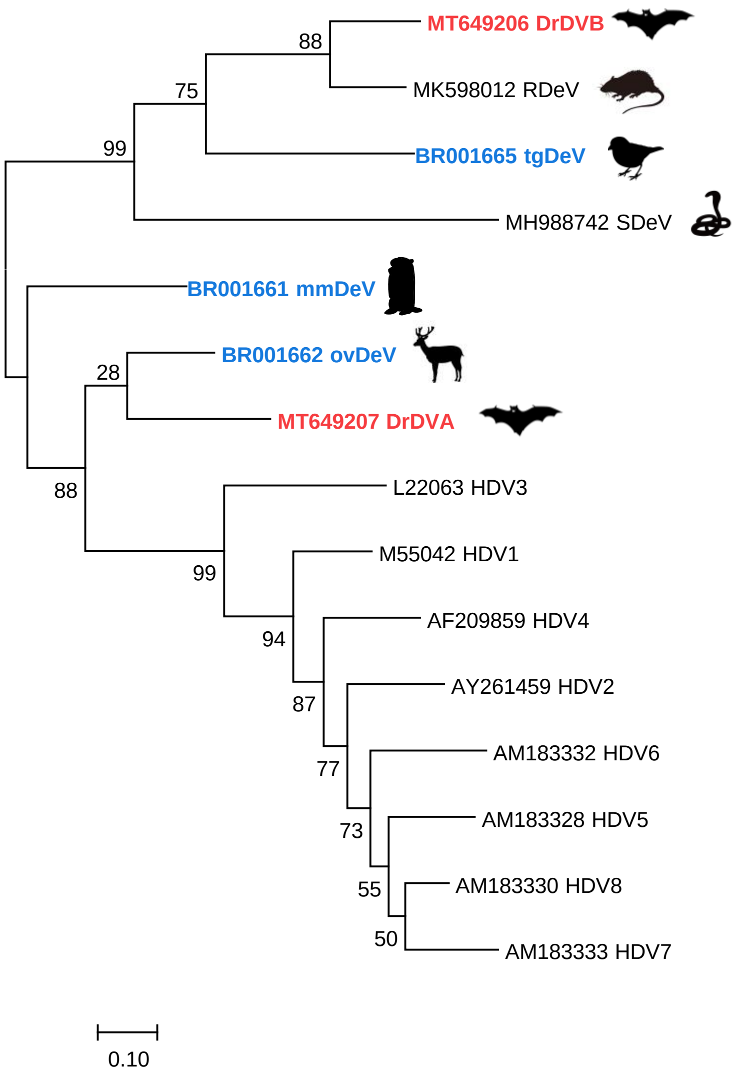


**Supplementary Figure 7. Phylogenetic relationship of deltaviruses.**
The phylogenetic tree was generated using the maximum likelihood method using an amino acid sequence alignment of representative deltaviruses. The deltaviruses identified in this study and bat deltaviruses are indicated in blue and red, respectively.

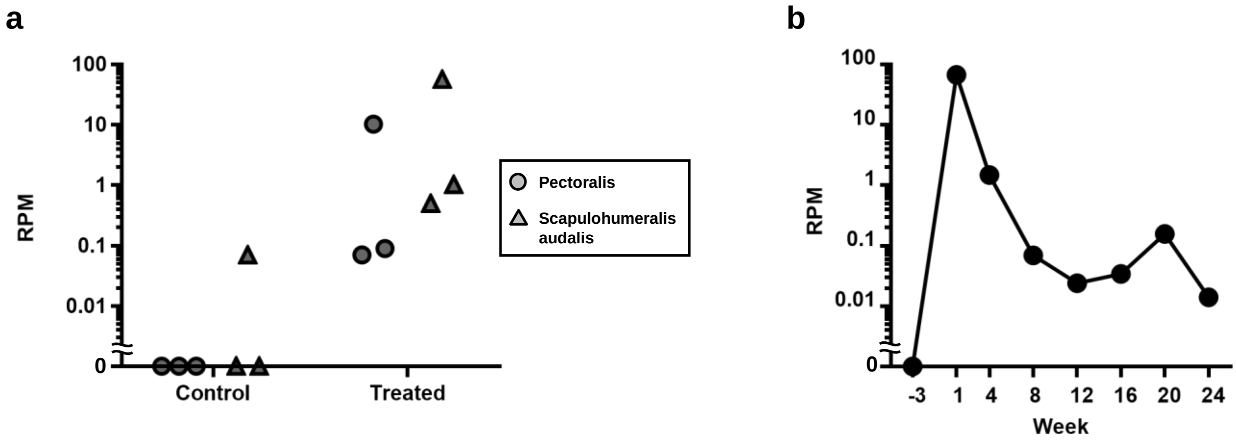


**Supplementary Figure 8. Newly identified deltaviruses might be susceptible to host's immune response.**
**(a)** Effects of testosterone treatment on tgDeV replication, transcription, or both. Mapped reads of the tgDeV genome in SRR2545941-SRR2545952 are shown as reads per million. The detailed data are shown in Supplementary Table 5. **(b)** Temporal transition of mmDeV-mapped reads in a woodchuck (SRR2136906-SRR2136918). The detailed data are shown in Supplementary Table 6.
