## Supplementary Table for "Identification of novel avian and mammalian deltaviruses provides new insights into deltavirus evolution"

**Supplementary Table 1. Summary of novel deltaviruses (full version)**

| Virus name | Host species | | Tiussue | SRA accession | DDBJ accession | Contig length (nt) | GC content (%) | BLASTx best hit | | |
|---|---|---|---|---|---|---|---|---|---|---|
| | Species name | Common name | | | | | | Virus name | Accession | Identity (%) |
| Taeniopygia guttata DeV | *Taeniopygia guttata* | Zebra finch | Scapulohumeralis caudalis | SRR2545946 | BR001665 | 1706 | 56.6 | Rodent deltavirus | QJD13558 | 63.3 |
| Marmota monax DeV | *Marmota monax* | Eastern woodchucks | Liver | SRR2136906 | BR001661 | 1712 | 53.4 | Hepatitis delta virus | AIR77039 | 60.0 |
| Odocoileus virginianus DeV | *Odocoileus virginianus* | White-tailed deer | Pedicle | SRR4256033 | BR001662 | 1690 | 56.4 | Hepatitis delta virus | AHB60712 | 66.7 |
| Erythrura gouldiae DeV | *Erythrura gouldiae* | Gouldian finch | Skin | SRR7504989 | BR001660 | 596 | 59.4 [a)] | Rodent deltavirus | QJD13555 | 63.5 |
| Serinus canaria-associated DeV | *Serinus canaria* | Domestic canary | Skin | SRR2915371 | BR001664 | 761 | 54.4 [a)] | Hepatitis delta virus | AIR77012 | 36.0 |

a) GC content of the partial genome sequences.

**Supplementary Table 2. Annotation of novel deltaviruses**

| Virus name | Position | | Annotation | Amino acids | Isoelectric point |
|---|---|---|---|---|---|
| | Start | End | | | |
| tgDeV | 1623 | 1030 | DAg ORF | 197 | 10.35 |
| | 969 | 964 | Poly(A) signal | | |
| | 689 | 773 | Genomic ribozyme | | |
| | 920 | 833 | Antigenomic ribozyme | | |
| mmDeV | 1629 | 1048 | DAg ORF | 193 | 10.40 |
| | 997 | 992 | Poly(A) signal | | |
| | 704 | 792 | Genomic ribozyme | | |
| | 946 | 858 | Antigenomic ribozyme | | |
| ovDeV | 1607 | 1023 | DAg ORF | 194 | 10.63 |
| | 973 | 968 | Poly(A) signal | | |
| | 701 | 783 | Genomic ribozyme | | |
| | 922 | 835 | Antigenomic ribozyme | | |
| scDeV | 756 | 139 | DAg ORF | 205 | 10.63 |
| | 90 | 85 | Poly(A) signal | | |
| egDeV | 574 | <1 [a)] | DAg ORF | >191 | 10.44 |

a) The ORF continues to outside of the obtained contig.

### Supplementary Table 3. Detection of deltavirus-derived reads in RNA-seq data (full version)

| Virus | BioProject/ BioStudy | SRA | Host | | | RPM (read per million) | Mappped read number | Total read number | Tissue |
|---|---|---|---|---|---|---|---|---|---|
| | | | Common name | Taxonomy | | | | | |
| | | | | Family | Species | | | | |
| tgDeV | PRJDB3398 | DRR087390 | Zebra finch | Estrildidae | *Taeniopygia guttata* | 0.02 | 2 | 93,969,226 | Brain |
| | PRJEB28085 | ERR2772428 | | | | 0.50 | 4 | 8,000,000 | Brain |
| | | ERR2772429 | | | | 0.50 | 4 | 8,000,000 | Brain |
| | | ERR2772431 | | | | 0.50 | 4 | 8,000,000 | Brain |
| | | ERR2772432 | | | | 0.25 | 2 | 8,000,000 | Brain |
| | PRJNA297576 | SRR2545941 | | | | 0.07 | 4 | 60,507,624 | Pectoralis |
| | | SRR2545942 | | | | 0.09 | 6 | 64,901,474 | Pectoralis |
| | | SRR2545943 | | | | 10.28 | 541 | 52,608,066 | Pectoralis |
| | | SRR2545944 | | | | 1.02 | 70 | 68,926,452 | Scapulohumeralis caudalis |
| | | SRR2545945 | | | | 0.50 | 30 | 60,435,950 | Scapulohumeralis caudalis |
| | | SRR2545946 | | | | 56.73 | 2,666 | 46,997,966 | Scapulohumeralis caudalis |
| | | SRR2545950 | | | | 0.07 | 4 | 53,458,024 | Scapulohumeralis caudalis |
| | PRJNA352507 | SRR5001843 | | | | 0.02 | 1 | 46,262,572 | Spleen |
| | | SRR5001847 | | | | 0.04 | 2 | 48,914,708 | Spleen |
| | | SRR5001848 | | | | 0.05 | 2 | 37,604,058 | Spleen |
| | | SRR5001849 | | | | 0.04 | 2 | 48,240,788 | Spleen |
| | | SRR5001850 | | | | 0.73 | 42 | 57,660,048 | Spleen |
| | | SRR5001851 | | | | 0.22 | 12 | 53,968,342 | Spleen |
| | PRJNA435424 | SRR6761870 | | | | 0.17 | 2 | 12,007,672 | Blood |
| | PRJNA413749 | SRR6151592 | Black-headed bunting | Emberizidae [a)] | *Emberiza melanocephala* | 0.11 | 2 | 18,837,970 | Liver |
| | PRJNA558524 | SRR9899549 | Black-headed bunting | | *Emberiza melanocephala* | 3.11 | 81 | 26,076,780 | Blood |
| | PRJNA470787 | SRR7244621 | Rufous-fronted bushtit | Aegithalidae | *Aegithalos iouschistos* | 0.03 | 2 | 78,465,516 | Liver |
| | | SRR7244624 | Rufous-fronted bushtit | | | 0.03 | 2 | 70,294,336 | Cardiac muscle |
| | | SRR7244629 | Yellow-bellied tit | Paridae | *Pardaliparus venustulus* | 0.01 | 1 | 72,167,958 | Lung |
| | | SRR7244650 | Grey crested tit | Paridae | *Lophophanes dichrous* | 0.03 | 2 | 79,286,956 | Liver |
| | | SRR7244673 | Rufous-vented tit | Paridae | *Periparus rubidiventris* | 0.03 | 2 | 62,012,430 | Lung |

| | | | | | | | | | |
|---|---|---|---|---|---|---|---|---|---|
| | | SRR7244693 | Yellow-bellied tit | Paridae | *Pardaliparus venustulus* | 10.68 | 666 | 62,346,176 | Lung |
| | | SRR7244695 | | | | 2.07 | 125 | 60,358,382 | Kidney |
| | | SRR7244696 | | | | 2.65 | 176 | 66,475,684 | Cardiac muscle |
| | | SRR7244697 | | | | 7.12 | 521 | 73,205,926 | Flight muscle |
| | | SRR7244698 | | | | 1.77 | 147 | 82,849,574 | Liver |
| | | SRR7244699 | Rufous-vented tit | | *Periparus rubidiventris* | 0.03 | 2 | 64,743,790 | Cardiac muscle |
| | | SRR7244728 | Black-throated bushtit | Aegithalidae | *Aegithalos concinnus* | 0.02 | 1 | 61,061,634 | Cardiac muscle |
| | PRJNA478907 | SRR7504989 | Gouldian finch | Estrildidae | *Erythrura gouldiae* | 1.07 | 51 | 47,450,142 | Skin |
| mmDeV | PRJNA291589 | SRR2136864 | Woodchuck | Sciuridae | *Marmota monax* | 0.04 | 3 | 79,395,232 | Liver |
| | | SRR2136865 | | | | 0.06 | 4 | 72,004,528 | Liver |
| | | SRR2136883 | | | | 0.07 | 4 | 60,179,086 | Liver |
| | | SRR2136906 | | | | 70.86 | 5,146 | 72,619,384 | Liver |
| | | SRR2136907 | | | | 63.08 | 4,358 | 69,088,520 | Liver |
| | | SRR2136908 | | | | 0.24 | 11 | 46,237,666 | Liver |
| | | SRR2136909 | | | | 0.05 | 2 | 40,860,302 | Liver |
| | | SRR2136910 | | | | 0.03 | 2 | 75,353,994 | Liver |
| | | SRR2136911 | | | | 0.04 | 2 | 47,486,488 | Liver |
| | | SRR2136912 | | | | 0.16 | 8 | 51,207,898 | Liver |
| | | SRR2136913 | | | | 0.03 | 2 | 69,102,274 | Liver |
| | | SRR2136916 | | | | 1.02 | 57 | 55,685,944 | Liver |
| | | SRR2136917 | | | | 0.90 | 38 | 42,002,154 | Liver |
| | | SRR2136918 | | | | 0.07 | 4 | 58,069,234 | Liver |
| | | SRR2136981 | | | | 0.04 | 2 | 47,345,042 | Liver |
| | | SRR2136982 | | | | 0.08 | 8 | 101,425,488 | Liver |
| | | SRR2136998 | | | | 0.04 | 2 | 47,208,236 | Liver |
| | | SRR2136999 | | | | 0.06 | 2 | 34,882,902 | Liver |
| | SRP011132 | SRR437934 | | | | 46.34 | 43 | 927,972 | PBMC |
| | | SRR437938 | | | | 19.83 | 22 | 1,109,424 | PBMC |
| ovDeV | PRJNA317745 | SRR4256026 | White-tailed deer | Cervidae | *Odocoileus virginianus* | 0.16 | 10 | 63,121,148 | Antler |
| | | SRR4256031 | | | | 0.18 | 12 | 66,438,392 | Muscle |
| | | SRR4256033 | | | | 180.73 | 8,456 | 46,787,594 | Pedicle |
| | | SRR4256034 | | | | 0.07 | 5 | 74,468,272 | Testis |
| scDeV | PRJNA300534 | SRR2915371 | Domestic canary | Fringillidae | *Serinus canaria* | 9.79 | 1,723 | 176,049,404 | Skin |

a) Emberizidae is regarded as the subfamily Emberizinae of the family Fringillidae in TimeTree.

#### Supplementary Table 4. Coexisting viruses in deltavirus-positive SRAs (full version)

| SRA | Host | | Query contig | | Subject | | | Query | | Subject | | E-value | Bitscore | Notes |
|---|---|---|---|---|---|---|---|---|---|---|---|---|---|---|
| accession | Species | Common name | Name | Length (nt) | Accession | Virus name | Identity | Start | End | Start | End | | | |
| SRR2545944 | *Taeniopygia guttata* | Zebra finch | NODE_79_length_906_cov_4.390832 | 906 | AUN86682 | Serinus canaria polyomavirus | 100.0 | 1 | 735 | 112 | 356 | 6.47E-178 | 505 | |
| SRR5001849 | *Taeniopygia guttata* | Zebra finch | NODE_709_length_1400_cov_37.345427 | 1400 | AUN86682 | Serinus canaria polyomavirus | 100.0 | 1349 | 282 | 1 | 356 | 0 | 726 | |
| | | | NODE_754_length_1362_cov_1.354086 | 1362 | AUN86683 | Serinus canaria polyomavirus | 96.2 | 1250 | 3 | 84 | 501 | 0 | 767 | |
| | | | NODE_1282_length_1068_cov_3.859738 | 1068 | AUN86680 | Serinus canaria polyomavirus | 100.0 | 82 | 1068 | 1 | 329 | 1.69E-167 | 481 | |
| SRR5001850 | *Taeniopygia guttata* | Zebra finch | NODE_348_length_1959_cov_5.914984 | 1959 | AUN86682 | Serinus canaria polyomavirus | 95.5 | 671 | 1738 | 1 | 356 | 0 | 677 | |
| SRR5001851 | *Taeniopygia guttata* | Zebra finch | NODE_297_length_1923_cov_5.157638 | 1923 | AUN86682 | Serinus canaria polyomavirus | 100.0 | 1261 | 194 | 1 | 356 | 0 | 726 | |
| SRR2915371 | *Serinus canaria* | Common canary | NODE_4_length_5248_cov_65.731580 | 5248 | YP_009041461 | Canary bornavirus 3 | 99.7 | 5248 | 125 | 6 | 1713 | 0 | 3300 | |
| | | | NODE_157_length_1443_cov_104.502196 | 1443 | YP_009041460 | Canary bornavirus 3 | 98.5 | 3 | 1379 | 43 | 501 | 0 | 909 | |
| | | | NODE_260_length_1228_cov_1158.091225 | 1228 | YP_009041458 | Canary bornavirus 3 | 100.0 | 1141 | 536 | 1 | 202 | 1.89E-127 | 375 | |
| | | | NODE_300_length_1177_cov_3498.696364 | 1177 | YP_009041456 | Canary bornavirus 3 | 100.0 | 41 | 1153 | 1 | 371 | 0 | 769 | |
| | | | NODE_1081_length_795_cov_9.143454 | 795 | NP_573443 | Canary circovirus | 96.7 | 705 | 79 | 42 | 250 | 7.02E-149 | 426 | |
| | | | NODE_1298_length_743_cov_2.166667 | 743 | NP_573442 | Canary circovirus | 97.9 | 743 | 36 | 55 | 290 | 2.90E-172 | 486 | |
| SRR7504989 | *Erythrura gouldiae* | Gouldian finch | NODE_114_length_1793_cov_16.827506 | 1793 | YP_009508817 | Erythrura gouldiae polyomavirus 1 | 95.3 | 1129 | 44 | 1 | 362 | 0 | 717 | |
| SRR7244695 | *Pardaliparus venustulus* | Yellow-bellied tit | NODE_2107_length_3092_cov_24.285332 | 3092 | ANE10779 | Mammalian rubulavirus 5 | 100 | 128 | 1654 | 1 | 509 | 0 | 1021 | Contamination? [a)] |
| | | | NODE_14943_length_1034_cov_35.282353 | 1043 | YP_009551871 | Rhinolophus gammaherpesvirus 1 | 33.673 | 704 | 997 | 16 | 111 | 6.98E-10 | 68.2 | Simple repeat [b)] |
| | | | NODE_15702_length_996_cov_3.342252 | 996 | AGT76392 | Human mastadenovirus C | 98.616 | 49 | 915 | 1 | 289 | 2.49E-174 | 494 | Contamination? [a)] |
| | | | NODE_3342_length_3230_cov_7.390929 | 3230 | ANE10783 | Mammalian rubulavirus 5 | 99.819 | 1670 | 18 | 1 | 551 | 0 | 951 | Contamination? [a)] |
| | | | NODE_7583_length_1982_cov_6.047265 | 1982 | ANE10784 | Mammalian rubulavirus 5 | 99.466 | 210 | 1895 | 4 | 565 | 0 | 1043 | Contamination? [a)] |
| | | | NODE_17287_length_870_cov_7.236057 | 870 | YP_009551871 | Rhinolophus gammaherpesvirus 1 | 33.673 | 555 | 262 | 16 | 111 | 4.83E-10 | 67.8 | Simple repeat [b)] |
| SRR7244697 | *Pardaliparus venustulus* | Yellow-bellied tit | NODE_21247_length_602_cov_5.924453 | 602 | YP_009551871 | Rhinolophus gammaherpesvirus 1 | 33.673 | 178 | 471 | 16 | 111 | 1.51E-10 | 67.8 | Simple repeat [b)] |
| SRR7244698 | *Pardaliparus venustulus* | Yellow-bellied tit | NODE_8399_length_1418_cov_4.649735 | 1418 | YP_009551871 | Rhinolophus gammaherpesvirus 1 | 33.673 | 316 | 609 | 16 | 111 | 1.37E-09 | 67.8 | Simple repeat [b)] |

a) These sequences showed 98-100% identities to human-derived viruses, and thus are highly probably derived from contamination or index hopping.

b) These sequences consist of direct simple repeats.

#### Supplementary Table 5. Primers and probes used in this study

| Name | Sequence (5'-3') | Note |
|---|---|---|
| tgDeV.F1 | CTTCGGTGGAATGGACTCTC | Real-time PCR detection of tgDeV from specimens |
| tgDeV.R1 | CAAGCGAAGAAGGCAAGAAG | |
| tgDeV.PCR.F1 | TTGGGACCCTTTTTCTTCGG | End-point PCR detection of tgDeV-like viruses |
| tgDeV.PCR.R1 | GACCCTTTCTCTGAGAGCAC | |
| tgDeV.PCR.F2 | TAACCCGAGAAAAGACCCAATGC | PCR and sequencing of lsDeV |
| tgDeV.PCR.R2 | AGGTGTCTTCAGCAATCCTCTCC | |
| tgDeV.PCR.F3 | CTTATAGGAGCGGCGGGGATAAC | PCR and sequencing of lsDeV |
| tgDeV.PCR.R3 | CTTAAGGGGAGGTTGGAACTCGG | |
| tgDeV.PCR.F4 | AAAGAGCAAGGAACTACGGAGGC | PCR and sequencing of lsDeV |
| tgDeV.PCR.R4 | GCTCTGATCCATTGACAGTTGCC | |
| tgDeV.PCR.F5 | TCGATTAACTTGCCCGTCTGTTG | PCR and sequencing of lsDeV |
| tgDeV.PCR.R5 | TGCTTACCCCGTAGATGTCTGTG | |
| tgDeV.PCR.F6 | CGAGGAGGTTCGAATGTCCGATG | PCR and sequencing of lsDeV |
| tgDeV.PCR.R6 | GCAAGCGAAGAAGGCAAGAAGC | |
| tgDeV.PCR.F7 | TTGGGAGTCGGGAACAAATCCTC | PCR and sequencing of lsDeV |
| tgDeV.PCR.R7 | GAAGGTGGAGACATGGATGTGGA | |
| tgDeV.PCR.F8 | GTCGGGACGATCCTCCATCTCC | PCR and sequencing of lsDeV |
| tgDeV.PCR.R8 | GGCCGACTCTGTCTTCTTTGAGA | |
| tgDeV.PCR.F9 | CGGGTTCGAGGCGAGCTACC | PCR and sequencing of lsDeV |
| tgDeV.PCR.R9 | CCGTGCTCGTCCACAAGGTAAG | |
| mmDeV.F1 | CTTCTTTCTCTAGCCTCATCCG | Real-time PCR detection of mmDeV from specimens |
| mmDeV.R1 | GAACATCCTGGGAATGGTGAG | |
| tgDeVAg-F | GAAACATTCCCTCCCCACGGGTG | Detection of tgDeV from the cells |
| tgDeVAg-R | GGACGCAACTCCGAAGAAAAAGGGTC | |
| mmDeVAg-F | GGAACTCCTGCTTTCCTCTAATG | Detection of mmDeV from the cells |
| mmDeVAg-R | GGAGAATCCTAAGCAAGGAAAC | |
| GAPDH-F | CCATGGAGAAGGCTGGGG | Detection of GAPDH from the cells |
| GAPDH-R | CAAAGTTGTCATGGATGACC | |
| human HDV-qF | GGACCCCTTCAGCGAACA | |
| human HDV-qR | CCTAGCATCTCCTCCTATCGCTAT | Quantification of human HDV from the cells |
| human HDV-qProbe | FAM-AGGCGCTTCGAGCGGTAGGAGTAAGA-TAMRA | |
| tgDeV-qF | GAAACATTCCCTCCCCACGGGTG | |
| tgDeV-qR | GACCCTTTTTCTTCGGAGTTGCGTCC | Quantification of tgDeV from the cells |
| tgDeV-qProbe | FAM- GGACGCAACTCCGAAGAAAAAGGGTC(MGB) | |
| mmDeV-qF | CTCTTTCTCACCATTCCCAGG | |
| mmDeV-qR | GGAGACTCTAAGACAATGGGTC | Quantification of mmDeV from the cells |
| mmDeV-qProbe | FAM- TTCCTCCTCGAGCTCCTCCTTTCT(MGB) | |
